# Predicting spread and effective control measures for African swine fever– should we blame the boars?

**DOI:** 10.1101/654160

**Authors:** Rachel A. Taylor, Tomasz Podgórski, Robin R. L. Simons, Sophie Ip, Paul Gale, Louise A. Kelly, Emma L. Snary

## Abstract

African swine fever (ASF) has been causing multiple outbreaks in Russia, Poland and the Baltic countries in recent years and is currently spreading westwards throughout Europe and eastwards into China, with cases occurring in wild boar and domestic pigs. Curtailing further spread of ASF requires full understanding of the transmission pathways of the disease. Wild boars have been implicated as a potential reservoir for the disease and one of the main modes of transmission within Europe. We developed a spatially explicit model to estimate the risk of infection with ASF in boar and pigs due to the natural movement of wild boar that is applicable across the whole of Europe. We demonstrate the model by using it to predict the probability that early cases of ASF in Poland were caused by wild boar dispersion. The risk of infection in 2015 is computed due to wild boar cases in Poland in 2014, compared against the reported cases in 2015 and then the procedure is repeated for 2015-2016. We find that long- and medium-distance spread of ASF (i.e. >30km) is very unlikely to have occurred due to boar dispersal, due in part to the generally short distances boar will travel (<20km on average). We also predict what the relative success of different control strategies would have been in 2015, if they were implemented in 2014. Results suggest that hunting of boar reduces the number of new cases, but a larger region is at risk of ASF compared to no control measure. Alternatively, introducing boar-proof fencing reduces the size of the region at risk in 2015, but not the total number of cases. Overall, our model suggests wild boar movement is only responsible for local transmission of disease, thus other pathways are more dominant in medium and long distance spread of the disease.

## Introduction

African swine fever (ASF) is a porcine disease that is rapidly spreading throughout Europe and Asia. Due to its high mortality rates and ability to spread within a farm, it can lead to significant costs for control and culling of large numbers of animals (Costard et al. 2013). The disease is a World Organisation of Animal Health (OIE) notifiable disease, thus identification of an outbreak can lead to international trade restrictions on pork products from that country. Combined trade restrictions and costly control measures indicate the potentially devastating economic impacts that are possible. The disease is endemic in sub-Sahara Africa, where it spreads chiefly between ticks and warthogs, bush pigs and wild pigs. However, an outbreak is currently ongoing in Asia and Europe, first identified in Georgia in 2007 followed by sporadic cases in Russia. In 2014, Lithuania and Poland were the first European countries to report and since then there has been steady spread westwards including in Latvia, Estonia, Hungary, Romania, Bulgaria, Ukraine, the Czech Republic and Belgium, by March 2019 (PAFF Committee 2019). Furthermore, in August 2018 the first reported cases of ASF in China occurred. By March 2019 there have been 114 outbreaks in 23 different provinces and spread to Mongolia and Vietnam (FAO 2019). Its arrival in China, with the world’s largest pork industry, highlights the seriousness of this disease for the global pig industry.

The disease appears to have a different transmission pattern in the recent outbreak in Europe and Asia compared to Africa, with the role of ticks inconclusive (Sánchez-Cordón et al. 2018). Many different transmission pathways have been implicated, and it is unclear which are more influential or if it is highly dependent on individual situations. In China, the majority of the cases appear to be related to the process of swill feeding (FAO 2019). Many of the cases on pig farms in the European outbreak have occurred in areas where cases of ASF-infected wild boar have also been found, especially in Latvia, Lithuania, Estonia and Poland, and hence wild boar contact has been implicated as one of the main modes of transmission (EFSA et al. 2015, Sánchez-Cordón et al. 2018). Natural movement of wild boar, including both home range movement and long range dispersal, could lead to contact of infected boar with susceptible boar and pigs in uninfected areas, thus spreading the disease away from the initial source. For transmission to pigs it is considered more likely that human-mediated transmission has been involved (Chenais et al. 2019) as direct contact between boar and pigs will be very low if sufficient biosecurity is in place. Long-distance jumps of the disease have also occurred, such as the cases in Czech Republic and Belgium which were several hundred kilometres away from the nearest case (EFSA et al. 2018), and these are very unlikely to have occurred due to natural wild boar movement. Human-mediated pathways are thought most likely to have led to those cases, with human transportation of infected meat perhaps most probable, although contaminated trucks, and movement of infected pigs, boar or boar carcasses are also possible transmission routes (EFSA et al. 2018, Chenais et al. 2019).

We focus on the role of wild boar in order to determine the extent to which natural movement of wild boar could be responsible for the spread of ASF at different distances. We create a model of wild boar movement and apply this to the early cases of ASF in Poland as a case study. The aim is to understand in detail how transmission of ASF due to wild boar movement can occur, in order to determine more accurately the risk of infection of ASF across Europe. Our model does not include other methods of transmission such as contaminated trucks, trade in pigs, or any human-mediated movement of boar or pig/boar meat, as our aim is not a comparison of risks due to different transmission pathways. However, our model is based upon a framework for estimating risk spatially for any disease and for any pathway (Taylor et al. 2019). Thus, the overarching model framework we present for this pathway is generic for any other disease spread by natural movement of wild terrestrial animals.

Other models have also considered the role of wild boar movement in transmission of ASF within Europe. For example, De la Torre et al. (2015) and Bosch et al. (2017) assess the risk of introduction of ASF due to wild boar at a country level for those countries free of ASF disease using a semi-quantitative method. Quantitative methods include those of Simons et al. (2019), which assesses risk of entry at a country level across Europe, using boar abundance and suitability at a fine scale in order to compute this risk, and the models of Thulke and Lange (2017) and Lange et al. (2018). In the latter two, boar movement is modelled on a fine spatial scale using individual-based modelling but on an abstraction, i.e. the spread of disease is not based on an actual European map but on a representative square grid. Alternatively, Iglesias et al. (2018) and Podgórski and Śmietanka (2018) use statistics to assess the role of wild boar movement in driving the outbreaks of ASF at various locations in Russia and Poland respectively. The former found that wild boar were responsible for 55% of new infections of ASF in their study area, whereas the latter found no significant correlation between statistics describing wild boar movement and new cases of ASF. In this paper, we take the model of Simons et al. (2019), but adapt it to assess risk of infection instead of introduction and on a finer spatial scale rather than only at a country level. Thus, this model is based upon actual European data for boar abundance as well as prevalence of cases, in order to assess risk of infection for specific locations in Europe, using Poland as a case study. We do not use an individual-based model like Thulke and Lange (2017) because we require our model to be generic to other terrestrial wild animals and due to the complexity it would involve on a large spatial scale, if expanding the model to assess risk across the whole of Europe.

The abundance of wild boar in Europe has been increasing over recent decades alongside an expanded range within Europe and colonisation of new habitats, including urban areas (Sáaez-Royuela and Telleriia 1986, Massei et al. 2015). This has prompted research on boar ecology, such as their movement, location and the suitability of habitat across Europe to sustain boar populations (Morelle et al. 2015). Wild boar exist in matrilineal groups, with separate home ranges for males and females and little interaction outside of the breeding period (D’Eath and Turner 2009, Podgórski et al. 2014). Males tend to have larger home ranges than females, with both varying seasonally. The main factor in long distance movement of boar is the post-weaning of young male boar, in search of new territory. This dispersal event usually involves a long distance movement outside of the natal home range for at least 40-50% of young male boar between 6 months and 2 years old (Truvé and Lemel 2003, Podgórski et al. 2014). Female boar may also choose to disperse, although usually for shorter distances than males (Podgórski et al. 2014). The combination of home range and dispersal is a significant factor in the difficulty in predicting the effects of the movement on disease spread by boar, especially since it is unclear how infection affects individual boar and their propensity to move longer distances. Although infected boar would still be able to disperse during their latent period, the virulence of the disease may reduce boar dispersal distance or the proportion of boar dispersing. Since wild boars are not subject to the same level of surveillance as commercial pig holdings, not only can they continue to have contact with other wild boar and potentially domestic pigs while alive and infected, but after death the carcass can remain in the environment for months (Probst et al. 2017). ASF virus is known to have a long survival time so can survive in the carcass and surrounding environment during this time (Mebus et al. 1993) and studies have shown it is not uncommon for wild animals, including other wild boar, to forage around dead carcasses (Probst et al. 2017). Thus, transmission from carcasses is a potentially important transmission route and we also consider its role in transmission within our model.

European countries and agencies are eager to know how best to control, manage and hopefully eradicate the disease, in order to stop the continued spread throughout Europe and the deleterious effects on the pig industry. A number of control strategies are implemented based on a European Union directive, such as culling, cleansing and disinfecting on infected farms, testing and removal of any boar carcasses found and increased awareness and promotion of biosecurity measures (European Union 2002). This directive also involves creating different zones around the infected cases – although the specific decision about how to manage the different zones is left up to the relevant country. In Belgium and the Czech Republic, after the first cases were discovered, a fence was built surrounding the infected area and no hunting or feeding of feral pigs was allowed within the area. Furthermore, all human activity in the area was stopped or reduced. In the second surveillance zone, hunting occurred to reduce the boar population outside the infected area. In both zones, increased passive surveillance was undertaken aiming to determine the extent of the disease and reduce the length of time boar carcasses may be in the environment. In both cases, this has been successful (so far) in preventing further local spread of the disease in Belgium and the Czech Republic (Mlynar 2018). The success, however, was undoubtedly only possible as the disease was found before it had led to a widespread outbreak.

To assess the effectiveness of the model we apply it to a Poland case study. In particular, we consider whether boar movement alone can be responsible for the cases that occurred in 2015 and 2016. We also consider the role of the three control strategies of carcass removal, hunting and fencing separately and we assess their effectiveness by comparing against the baseline predicted risk in 2015 without the control strategies.

## Methods

We create a model of wild boar movement in order to calculate the risk of initial infection with ASF in new areas. We include various factors related to ASF epidemiology and wild boar ecology such as transmission by live boar and by carcasses as well as the different movement behaviours of boar at different life stages. We outline the overall generic risk assessment model and then how we adapted this to the specific case study of wild boar terrestrial movement. We use the early cases of ASF in Poland, from 2014-2016, as a case study because the initial cases of ASF in Poland were all in wild boar and in a localised area. The first cases were all in close proximity to the border with Belarus, where 2 cases had been reported in 2013, suggesting transboundary introduction as the initial source (Pejsak et al. 2014). We use the reported cases of ASF in 2014 in Poland to predict the probability, due to wild boar movement only, of at least one infection in 2015 in the area surrounding the 2014 cases including the case locations. We perform the same method to predict from reported cases of 2015 the probability of at least one infection in 2016. In both cases, we compare our predicted probability of at least one infection in 2015/2016 against the locations of reported cases in 2015/2016. We then consider various realistic scenarios of the potential control measures of hunting, fencing and increasing the removal rate of carcasses, and assess their effectiveness by comparing against the baseline scenario for risk of infection in 2015.

### Overview of the risk assessment model

We model the risk of infection of ASF within new areas in Europe due to wild boar movement using the spatial quantitative risk assessment framework outlined in Taylor et al. (2019). This framework incorporates many disease entry pathways and for each considers the number of infectious animals (or infectious units depending on the entry pathway) entering a new Area B given that the disease is present in Area A, whether detection of these animals would occur, survival of these animals in the new Area B, and contact and transmission with susceptible animals in Area B. Adopting this framework, we calculate the risk of initial infection with ASF using a cell structure across Europe where each cell is 100km^2^, i.e. with sides of 10km. Area A is defined as those cells which we estimate to have non-zero prevalence (see the section on prevalence below for how we calculate this). Area B is any cell within Europe (regardless of whether cases have already been reported in that cell). Following Taylor et al. (2019), the number of infected wild boar reaching cell *c* of Area B is given by:

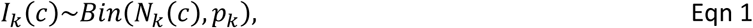

where *k* represents a cell in Area A, *N*_*k*_(*c*) is the number of wild boar moving from cell *k* to cell *c* and *p*_*k*_ is the prevalence in cell *k*. We thus define *I*(*c*) = ∑_*k*_ *I*_*k*_(*c*) as the total number of infected wild boar entering cell *c* from any cell *k*. We assume no detection of infection in wild boar will occur during their movement from one cell to another.

We model the contact with susceptible boars and pigs using the disease metric *R*_0_, which is a measure of the expected number of infections that would occur if one infected wild boar were to enter a susceptible population. Our equation for *R*_0_(*c*) includes information on the susceptible populations at risk in cell *c*, the pathway of entry, the transmission between infected and susceptible animals and the survival of infection. Each parameter from *R*_0_(*c*) is drawn from a distribution of likely values. The number of initial infections, *N*_*I*_(*c*), in cell *c* is then calculated by *I*(*c*) draws from a Poisson distribution with mean given by *R*_0_(*c*):

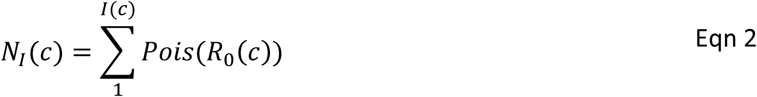

That is, there are *I*(*c*) infected wild boar entering cell *c*, each of which has a mean probability of infecting *R*_0_(*c*) susceptible animals, to give *N*_*I*_(*c*) new infections. The probability, *R*_*I*_(*c*), that at least one infection in a susceptible animal would occur in cell *c* is then given by the proportion of the simulations where infection occurs in a susceptible boar or pig. We outline in greater detail below how we compute each step of the risk assessment for the specific wild boar movement pathway.

### The wild boar movement model

We model the number of boar moving from Area A to Area B based on wild boar movement ecology, using the cell structure mentioned, on a yearly timescale. We adapt a model of Simons et al. (2019), which models the number of infected wild boar which will reach the border of a new country. However, we enhance the model to estimate at a cell level where the boar are likely to travel to, regardless of country borders. For each cell in Area A, we estimate where those boar will be after a year, depending on whether they perform one or two types of movement, namely home range movement and long range dispersal. All boar have a home range. Long range dispersal may be undertaken by adult male and female boar but is predominantly undertaken by young male adult boar, looking for new territory. For long distance movement, we assume the direction the boar moves is determined by the habitat suitability of each cell, where habitat suitability is a measure of how suited each cell is for boar to live and is given by a score between 0 and 1. Thus, boar movement is a biased random walk, characterised by the benefit they will receive from moving to the neighbouring cells, to represent the fact that boar are moving in order to find better territory (Truvé and Lemel 2003). Given the long range distance that boar travel, we a fix a total number of steps (defined as a movement from one cell to a neighbouring cell) the boar can travel, *n*, to be the total distance in km divided by the width of a cell, in our case 10km. We use *N*(*c*) to indicate the neighbourhood of each cell *c*, specifically the 8 cells to the north, east, south and west, as well as the cell itself. Thus boar can choose to stay in their current cell if it has a high suitability. The probability that a boar will be in cell *c* after *n* steps, *p*(*c, k, n*), given that the boar started from cell *k* in Area A, is then given by the recurrence relation:

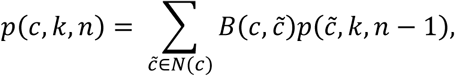

where 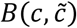 is a probability of moving from cell 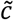 to cell *c* determined by the benefit a boar will receive by moving, and is multiplied by the probability of being in cell 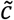 after *n* − 1 steps. The recurrence relation has initial conditions at step 0 of:

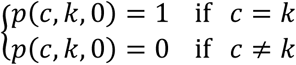

Note that for cells neighbouring *k*, the origin cell, we do not allow movement back to *k* in order to ensure dispersal from the home range does actually take place. The benefit probability 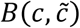 is based on the habitat suitability score, *h*(*c*), in each cell *c*. It is calculated by comparing the difference in the suitability between the two cells *c* and 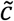 and normalising this by the benefit that could be gained by moving to any of the neighbouring cells *c*\* of cell 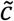:

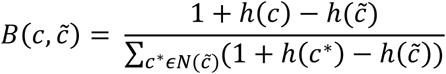

Since most boar dispersal is by young adult boar who perform the long-range movement in search of new territory usually only once in their lifetime, we model one movement event of boar and then assess the risk for a year, but do not specifically model the time that events occur within the year. Once boar arrive at their final destination, we assume they perform home range movement in this new cell for the rest of the year. We also include a probability (*p*_*i*_) that the boar will be infected at the same time as their long distance dispersal.

For home-range movement, we assume that the boar can travel anywhere within the home-range area regardless of the habitat suitability. In practice, the home-range is smaller than the cell size and therefore we assume that the boar can explore a percentage of the cell during home-range movement, dependent on the home-range size.

### Abundance and suitability maps

We use wild boar abundance and habitat suitability maps from Alexander et al. (2016). To create the habitat suitability map, Alexander and co-authors used land cover databases, published descriptions of boar preferences and expert opinions. Combining the habitat suitability map with abundance-related data, from sources such as national and international databases and hunting records, within a species distribution model, they produced a European wide estimate of boar abundance. They outputted the boar abundance estimates on a semi-quantitative scale of 0-4 for each cell, rather than as specific values. We reconstructed a numerical value for boar density per cell by manually curating the papers cited by Alexander et al. (2016), and extracting the data values from these papers to calculate population densities. We then calculated the quartiles of our population estimates so that we could substitute the semi-quantitative scores of Alexander et al. (2016) with the quartile values instead. For both maps, Alexander et al. (2016) produce output on a 1km^2^ resolution. However, we aggregate the cell resolution up to 100km^2^ in order to speed up calculations within the movement model.

Our model calculates the risk of initial infection of ASF occurring in a susceptible population, whether that population is wild boar or pigs. Therefore we need abundance maps of the susceptible animal populations as well. For the boar population, we use the same boar abundance maps as above, but we reduce the number of animals in each cell by the number of estimated cases of ASF in boar in that cell in order to get a total number of susceptible boar. For the pig density maps, we use data from the FAO gridded livestock of the world (FAO 2014).

### Prevalence in wild boar

We estimate the prevalence of ASF in wild boar across Europe by extracting data on the number of cases of ASF from Empres-i, including case locations. Empres-i gathers its information on outbreaks from multiple sources, although the major source is the OIE. Other sources include FAO officers, European Commission and media. We aggregate the total number of wild boar cases by each 100km^2^ cell and then multiply this total by an under-reporting factor. Since all boar hunted or found dead in Poland are tested for ASF, this underreporting factor is predominantly comprised of the likelihood of finding carcasses. We also use this value to determine the carcass removal rate in the baseline scenario. We then calculate a first estimate of the prevalence by dividing the inflated (by underreporting) number of cases by the number of boar in that cell. However, it is possible that cases have occurred in neighbouring cells and also have not been found, correctly diagnosed or reported, since boar move irrespective of our cell boundaries. We therefore perform a smoothing method to the prevalence data to account for this, which spreads our inflated cases across neighbouring cells. It does this by recalculating the prevalence in all cells as a weighted average between the current prevalence in that cell and the prevalence in all 8 neighbours, with 50% weight given to the current cell and 50% weight distributed equally among the neighbouring cells. We then recalculate the estimated number of cases in each cell by multiplying the number of boar by the new prevalence estimate. Smoothing the prevalence and then recalculating the number of cases (rather than smoothing the number of cases) ensures that we take into account the abundance of boar and hence the fact that some cells may have no cases due to no boar being present in that cell rather than underreporting. Estimated prevalence based on the cases in Poland in 2014 is plotted in Figure 1.

**Figure 1.**
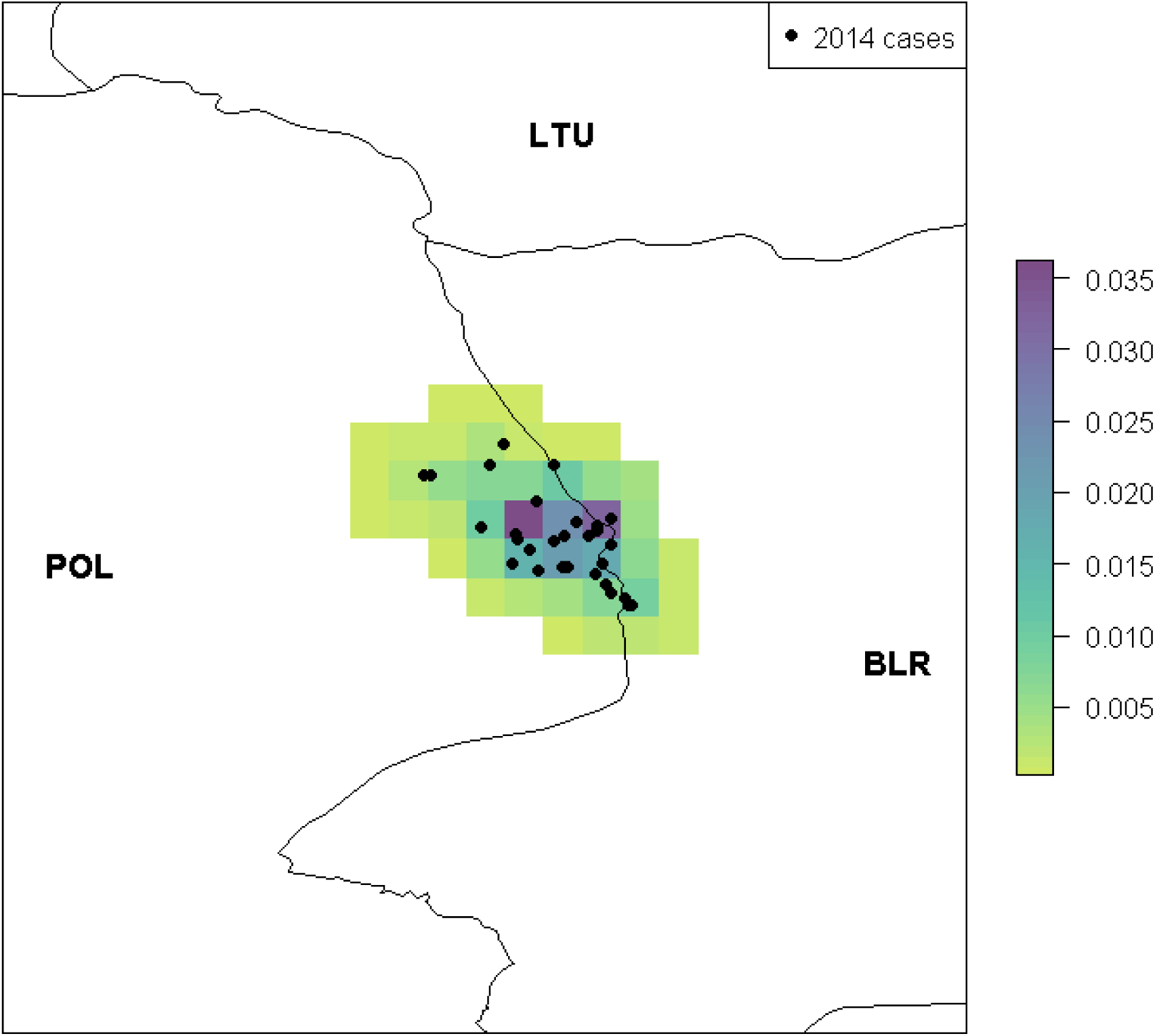
The estimated prevalence at a 100km^2^ cell level, given the reported cases in Poland in 2014 and the boar abundance. Prevalence is represented at a cell level by the colour scale on the right. Black circles indicate the locations of reported cases (which may have had >1 infected wild boar). White indicates zero prevalence. Countries are indicated by their ISO3 code.

### Contact and transmission

We create two separate equations for *R*_0_(*c*) in order to separately assess contact and transmission with live pigs and with live boar. We model the contact with susceptible pigs within each cell *c* by considering how many susceptible animals are in the cell (*S*(*c*)), the length of the infectious period (1/*r*) and the per-capita contact rate (*γ*) and transmission probability (*β*) between species:

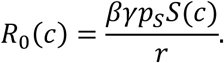

We multiply *S*(*c*) by *p*_*S*_, the proportion of susceptible animals in the cell that the infected animal is likely to be in contact with, because the home range is smaller than our cell size (100km^2^) and thus the boar will not roam the whole cell but only a proportion of it. Therefore *p*_*S*_ is the proportional size of the home range compared to the cell size.

Wild boar predominantly exist in matrilineal groups (Podgórski et al. 2014). Therefore, for boar contact with other live boar we use the equation as above but modify it to include both within-group contact and between-group contact, by replacing *γp*_*S*_*S*(*c*) with

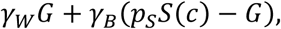

In this equation, *γ*_*W*_ is the within-group contact rate, *G* is the average group size and *γ*_*B*_ is the between-group contact rate which is applicable for contact with all other boar in the home range. We assume that only a proportion of boar survive infection. The carcasses of boar that die due to ASF can also contribute to transmission of ASF. For these carcasses, we estimate how many live boar will have direct contact with the carcass and thus able to become infectious by using a study of Probst et al. (2017), in which data on live boar contact with boar carcasses is collected. We compute the number of new cases of ASF due to carcasses as:

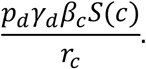

Here, *p*_*d*_ is the probability that direct contact will occur with a carcass (direct contact does not always occur due to the time of the year of death, location of carcass etc.), *γ*_*d*_ is the total number of direct contacts per year each boar has with a carcass, *β*_*c*_ is the transmission probability from a carcass to a susceptible animal per contact, *r*_*c*_ is the rate at which the carcass is available to cause infection, which is the inverse of the length of time the carcass is available (*T*_*c*_). This is determined by two factors, skeletonisation of the carcass and whether the carcass is removed, as follows:

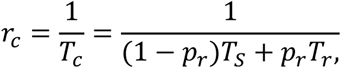

where *T*_*S*_ is the time until skeletonisation of the carcass, *T*_*r*_ is the time until removal of a carcass and *p*_*r*_ is the probability that a boar carcass is found and removed.

Therefore, our full equation for *R*_0_(*c*) in susceptible boar populations, i.e. the likelihood of new cases occurring in susceptible boar in cell *c* given an infected wild boar has entered the cell, is:

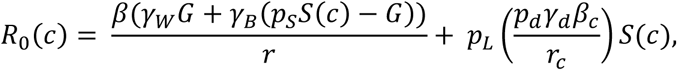

where *p*_*L*_ is the probability of lethal infection in boar.

We then calculate the number of new infections in boar or pigs occurring in cell *c* because of infected animals *I*(*c*) moving in to cell *c* by eqn 2. For our results, firstly we calculate quantiles of the number of new infections in boar or pigs in each cell. Secondly, the overall probability of at least one infection in each cell is calculated based upon the proportion of simulations in which infection occurs in that cell. Lastly, we define the risk region as all of the cells which have non-zero probability of at least one infection.

### Parameter Values

A full list of all parameter values, their descriptions and references is provided in Table 1. We outline the choice for key parameters.

**Table 1.**
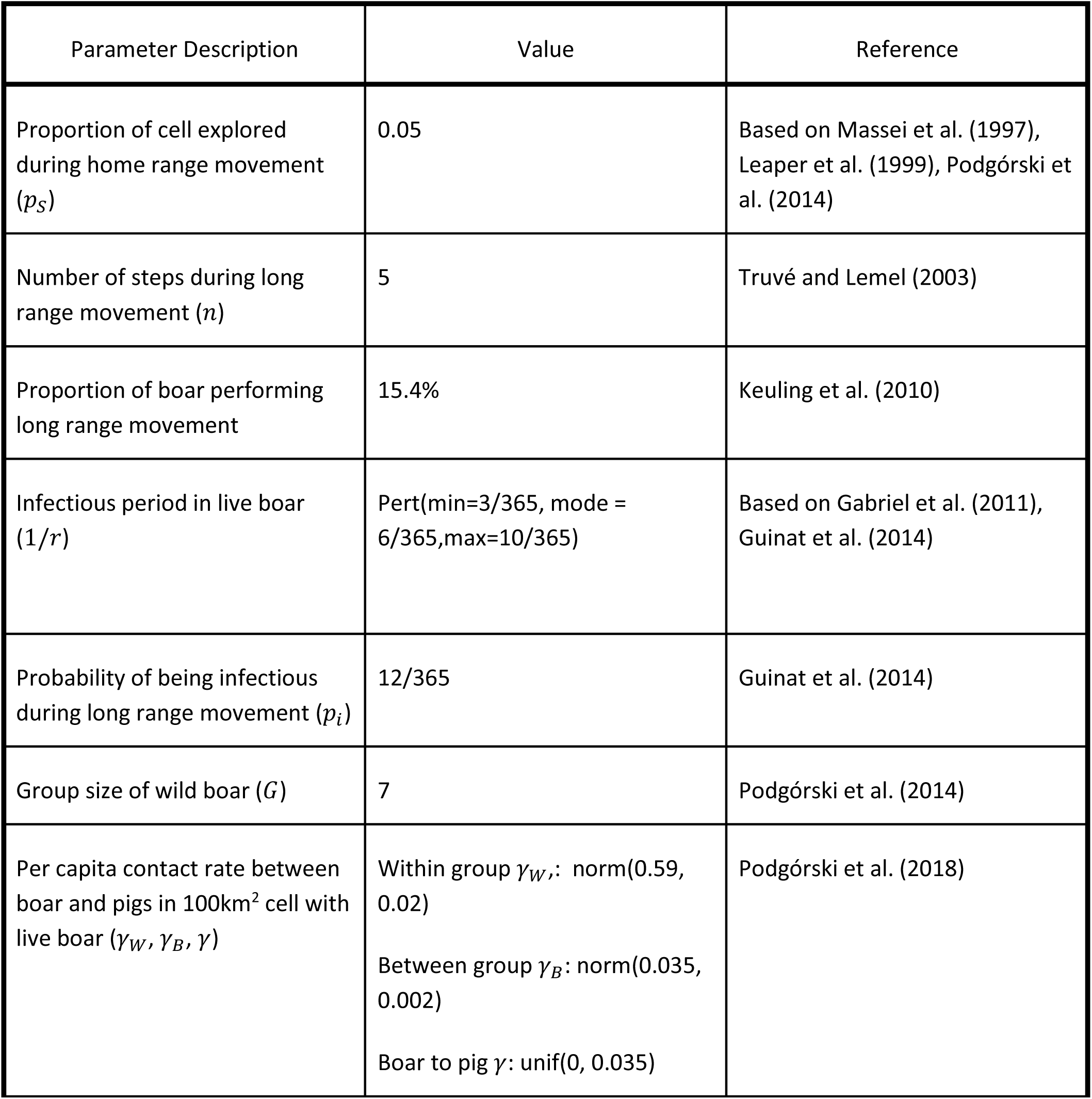
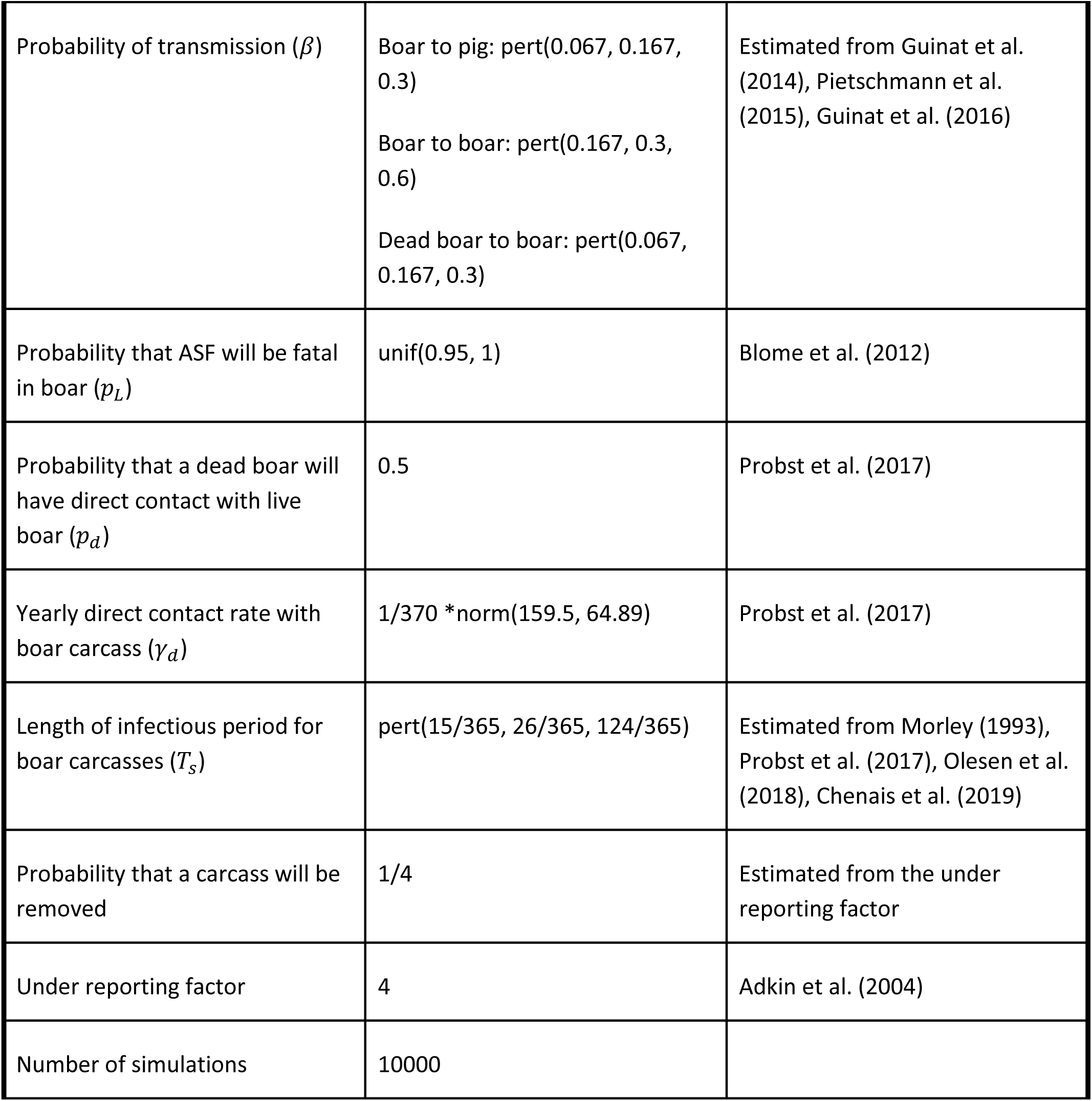
Parameter values with a description and reference. Rates and times are given in units of years unless specified otherwise.

In order to calculate the proportion of the cell explored during home range movement (*p*_*S*_), we used an average home range size of 5km^2^ based upon various studies (Massei et al. 1997, Leaper et al. 1999, Podgórski et al. 2014). For long distance movement, the model considers how many steps the boar will take based on how many kilometres they travel. Therefore, the parameter is defined as total distance travelled by boar rather than the straight-line distance from their starting point. Estimates for dispersal distance of boar are wide-ranging, such as less than 30km for most boar but up to 100km possible (Truvé and Lemel 2003), a maximum dispersal distance of 41.53km (Keuling et al. 2010), 48km (Lange 2015) and even up to 250km (Andrzejewski and Jezierski 1978). However, these estimates are all based on straight-line distances from original location. Therefore, we choose 50km as the dispersal distance as this allows for longer movement but most boar will disperse less than 50km from their original location similar to Truvé and Lemel (2003). Furthermore, it incorporates the fact that a boar with infection of ASF is unlikely to perform a very long movement. Thus the number of steps a boar can take, *n*, is 5. We calculate the probability that a boar will be infected when they start their dispersal event (*p*_*i*_) by dividing the latent period for indirect contact, estimated at 12 days (Guinat et al. 2014), by 365.

We calculate the contact rates between live boar and other live boar using Podgórski et al. (2018) which estimated within-group and between-group contacts rates to be 0.59 ± 0.02 and 0.035 ± 0.02 respectively. We use the mean of between-group contact (0.035) as an upper limit in a uniform distribution for boar contact with pigs, with a lower limit of 0, to represent uncertainty in this parameter based on biosecurity of farms, but that it wouldn’t be higher than contact with other boar.

For contact between live boars and boar carcasses, we primarily used the study of Probst et al. (2017), which used cameras to assess the level of contact between live boar and boar carcasses over the course of a year. Of 32 carcasses laid out, 16 experienced direct contact (thus *p*_*d*_ = 0.5). Of those 16 carcasses, 189 direct contact visits by live boar were counted, an average of 11.8 visits per carcass. We assume there is a normal distribution around this estimate with a standard deviation of 4.8 around 11.8 based on the average number of visits per carcass site. However, due to skeletonization of the carcass, each carcass was approximately available for direct contact for 3.8 weeks (27 days) only. Therefore, our yearly contact rate (shown using the mean of carcass visits, 11.8) is 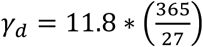 where 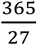 converts the contact rate from a rate per 27 days to a rate per year. However, we convert this into a per boar contact rate by estimating the mean number of boar that were in the area where the carcasses were placed – which we estimate to be 370. Therefore the contact rate per boar is calculated by dividing *γ*_*d*_ by 370.

The length of the infectious period for carcasses (*T*_*c*_) is also determined by the time until skeletonization (*T*_*s*_). As ASF virus has been found to survive in meat for up to 230 days (Morley 1993), we assume that an infected carcass will be infectious until skeletonization is complete. However, we adjust the length of the infectious period to incorporate two factors – possible transmission after skeletonization through contact with both the soil surrounding the remaining bones and chewing the bones themselves, and the fact that boar are unlikely to contact a boar carcass within the first 15 days (Probst et al. 2017). Based on a time until skeletonisation of *T*_*s*_ = 27 (Probst et al. 2017), but lack of boar contact for the first 15 days, we set 12 days to represent the average time before skeletonisation during which boar may contact the carcass. Regarding the length of time in which boar may contact the remaining bones or soil below the carcass, there is no consensus over how long the virus could remain in the environment or bones. In one study environmental transmission due to shedding of virus from infected live boar was found to be less than 3 days (Olesen et al. 2018). Chenais et al. (2019) state that ASF has been found experimentally to remain infectious in forest soil for 112 days. Further, Chenais et al. (2019) mentioned another study in which soil samples from carcass locations were PCR-positive “several days or weeks after the carcasses has been removed although no viable virus could be isolated”. We therefore use a pert distribution of pert(3,14,112) days to describe the length of time that transmission is possible after skeletonisation.

We also include the probability of a boar carcass being found and removed (*p*_*r*_) and the length of time until this occurs (*T*_*r*_), to complete the calculation of *T*_*c*_. For the probability of the boar carcass being found, we assume that all reported boar cases were removed, and hence we use the inverse of the under-reporting factor in number of cases to estimate what proportion of the true cases are found and removed. The length of time until carcass removal is given by a uniform distribution from 1 day until the length of time until carcass skeletonisation.

For the probability of transmission for a direct contact between a live boar and a boar carcass, we use a pert distribution with mode of 0.167 due to a study by Pietschmann et al. (2015) in which boar are infected with a low dose of ASF corresponding to “those obtained through contact with fomites, swill, excretions of infected animals, or contact with carcasses.” Of the 12 boar that were inoculated, 2 became infected. Guinat et al. (2016) estimated the transmission rate for indirect contact to be 0.3 so we use this as the max of the pert. However, Guinat et al. (2014) also found that indirect transmission occurred within 6 to 15 days post infection, therefore we take 1/15 as the min of our pert distribution. For the probability of transmission between live boar and pigs, we assume the same distribution as for boar carcasses. For transmission between live boar with other boar, we assume the same min of the pert distribution but increase the mode to 0.3 and the maximum to 0.6, as Guinat et al. (2016) produce this latter estimate for transmission between boar in direct contact under experimental settings.

### Scenario Analyses

We perform different scenario analyses to address the potential control strategies that could have been implemented in Poland to assess their effectiveness when wild boar alone is responsible for transmission. We assume that the control strategy is implemented in 2014 and compute a new risk for 2015 including the potentially positive and negative effects of the control strategy. We then compare our new results against the baseline results for 2015.

The scenario analyses we perform are:

A. Increase in carcass removal (CR): This affects the probability of a carcass being found and removed; in the baseline model 1 in 4 carcasses are removed CR1. 1 in 3 carcasses are found and removed CR2. 1 in 2 carcasses are found and removed
B. Hunting (H): This affects both the number of boar in the area and the probability that boar will move due to disturbance H1. Reduction of boar population to half original size and 25% of remaining boars disperse H2. Reduction of boar population to quarter of original size and 25% of remaining boars disperse H3. Reduction of boar population to half original size and 50% of remaining boars disperse H4. Reduction of boar population to quarter of original size and 50% of remaining boars disperse
C. Fencing (F): The implementation of fencing around the current cases is determined by a buffer width and the permeability of the fence in successfully stopping boar moving outside the buffer F1. Fence of 10km buffer and 95% successful F2. Fence of 10km buffer and 50% successful F3. Fence of 20km buffer and 95% successful F4. Fence of 20km buffer and 50% successful

Please see Appendix A in the Supplementary Information for an explanation of how the scenario analysis were implemented in the model.

We assess the effectiveness of these 10 control scenarios by comparing them against the baseline scenario using three metrics assessing severity and spread, namely the total number of cases per simulation; the total number of cells which have cases per simulation; and the total number of cells which have a non-zero probability of at least 1 case over the 10,000 simulations. Thus, the first two metrics indicate what a single simulation looks like, regarding severity and spread, and how much this varies across all the simulations. The third metric is a summary statistic indicating overall potential spread by stating the number of cells that have a case in at least one simulation (i.e. the size of the risk region).

## Results

We plot the spatial probability of at least one infection of ASF in 2015 due to our estimated prevalence of ASF in 2014 in Figure 2. This is assuming that one dispersal event takes place each year. The risk to wild boars is significantly higher than the risk to pigs, with the highest probability in any cell being 0.025 for pigs, i.e. only a 2.5% chance that a new pig case would occur in that cell. The outer edges of the risk region have very low probabilities. As we are running 10,000 simulations only, it is possible that neighbouring cells to the edge could also have a non-zero probability of infection, which is less than 1×10^-4^. However, cells much further than one neighbour away from the risk region would have negligible risk according to our model due to our imposed limit for distance that boar can travel during dispersal.

**Figure 2.**
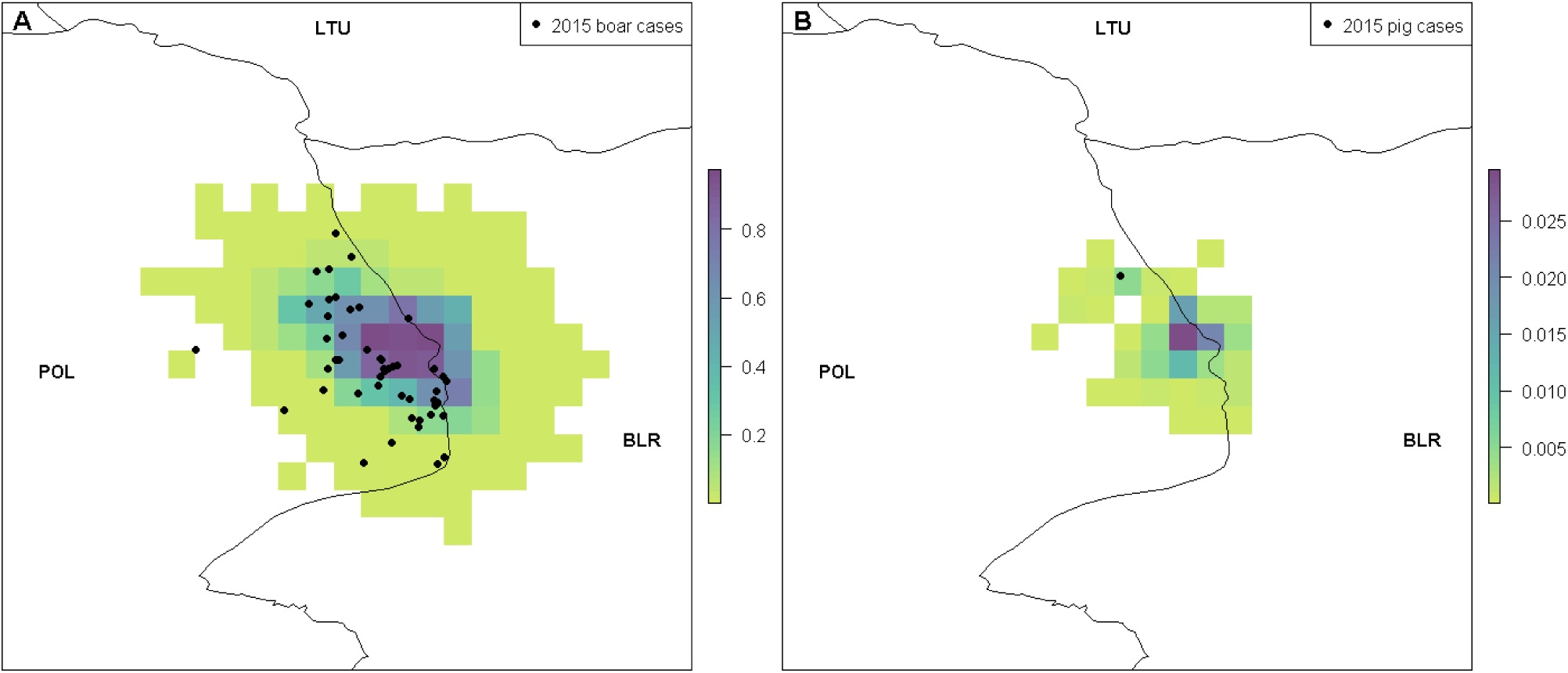
The probability of at least one infection in boar (A) and pigs (B) in 2015 caused by the movement of wild boar over one year. Black circles represent reported cases of ASF in boar and pigs in 2015. Countries are indicated by their ISO3 code.

Infection in boar could occur due to transmission from an infectious live boar or from contact with an infected carcass. The 2.5%, 50% and 97.5% quantiles of the number of new infections in boar are estimated separately for those caused by live boar contact or contact with carcasses (Figure 3). This highlights that infection in wild boar is predominantly caused by contact with carcasses (see also Appendix B for the probability of at least one case in 2015 in boar split by live boar or boar carcasses). The median and upper percentiles for live boar (Figure 3B, C) indicate lower numbers of infections compared to those due to carcass transmission (Figure 3E, F). This is because the estimate for *R*_0_ is higher on average for boar carcasses than contact with live boars since boar carcasses are in the environment and infectious much longer than the average length of time a live boar is infectious. For the lower percentile, no cases occur due to transmission by boar carcasses or live boar (Figure 3A, D), indicating that with a low probability it is possible that the outbreak in Poland in 2014 would die out, if only transmission by boar is responsible for spread. The upper percentile plots outline a smaller region of risk than the risk maps in Figure 2, indicating that in only a small percentage of simulations do we estimate that boar spread could be wider than this reduced region. In general, the number of cases that we predict is lower than the total number of boar cases that were reported (with an underreporting factor included). This is reasonable since our model is only predicting those first initial cases that occur due to boar movement and does not include any secondary cases which could happen subsequently in that new area within the course of the year.

**Figure 3.**
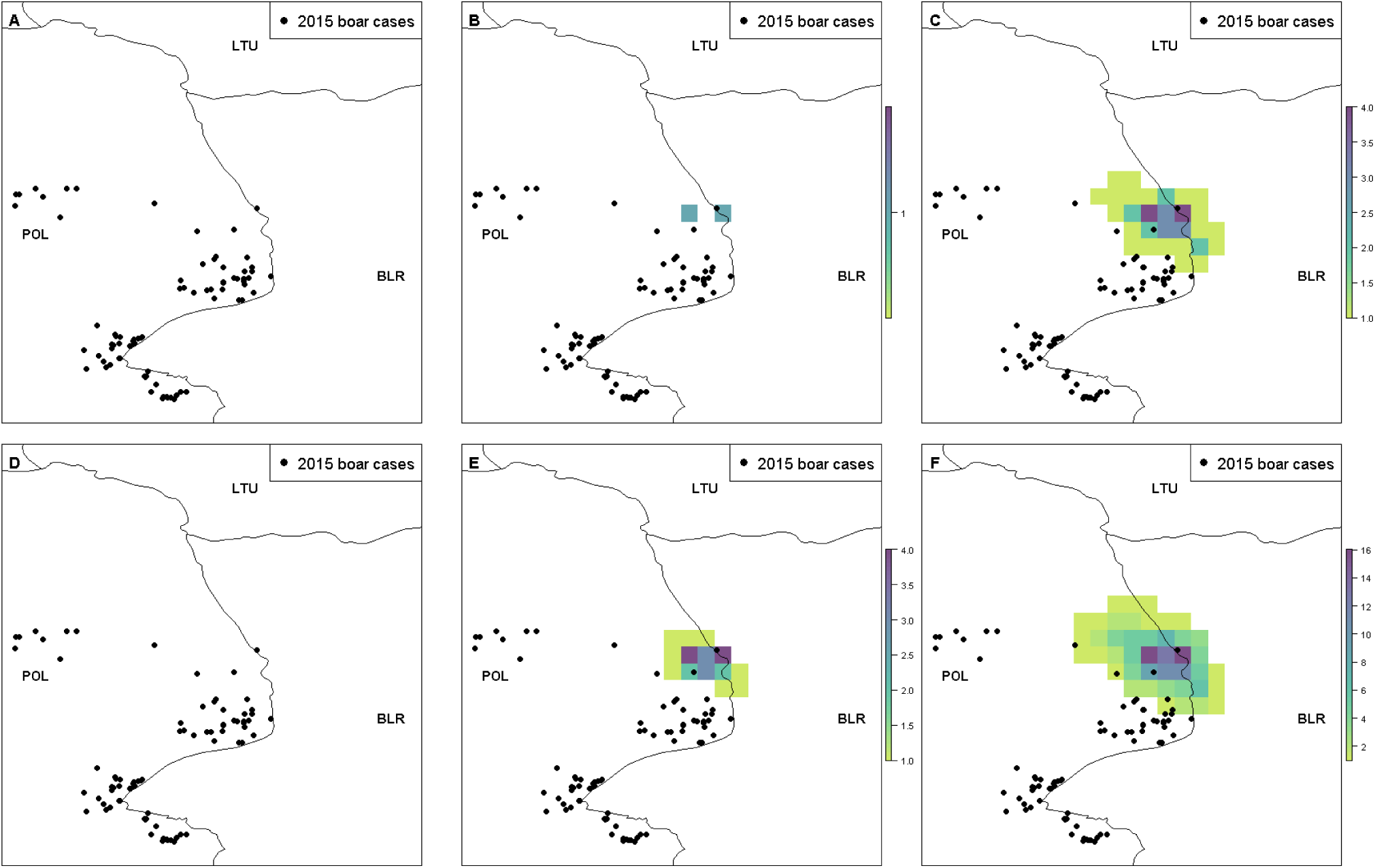
The 2.5%, 50% and 97.5% quantiles of the number of new infections in boar in 2015 due to transmission by live boar (A, B, C respectively) and by contact with boar carcasses (D, E, F respectively). Black circles indicate reported cases of boar in 2015. Countries are indicated by their ISO3 code.

The midpoint of the 2015 cases was 7.204km away from the midpoint of the 2014 cases and all of the cases in 2015 were within 9.163km of their nearest 2014 cases. Thus, the disease spread remained relatively confined in this year of the outbreak.

### Probability of new infections in 2016 given 2015 reported cases

We now predict the probability of at least one new infection of ASF in wild boar and pigs in 2016 due to movement of wild boar (Figure 4). The probability of ASF in boar is on a similar scale to that predicted for 2015, with over 80% chance of ASF infection in some cells. In comparison, the probability of ASF in pigs is much lower in 2016 than in 2015 with a maximum of 0.7% chance of transmission in a cell. However, in both cases the number of cells in which a non-zero probability of transmission occurs is much higher due to the movement of boar further out from the original cases in 2014 after two dispersal events. In Figure 4C and D, we plot separately the probability of cases occurring due to wild boar movement and boar carcasses. As before, this indicates that under the current parameterisation of the model, wild boar transmission is largely driven by contact with carcasses rather than with other live boar.

**Figure 4.**
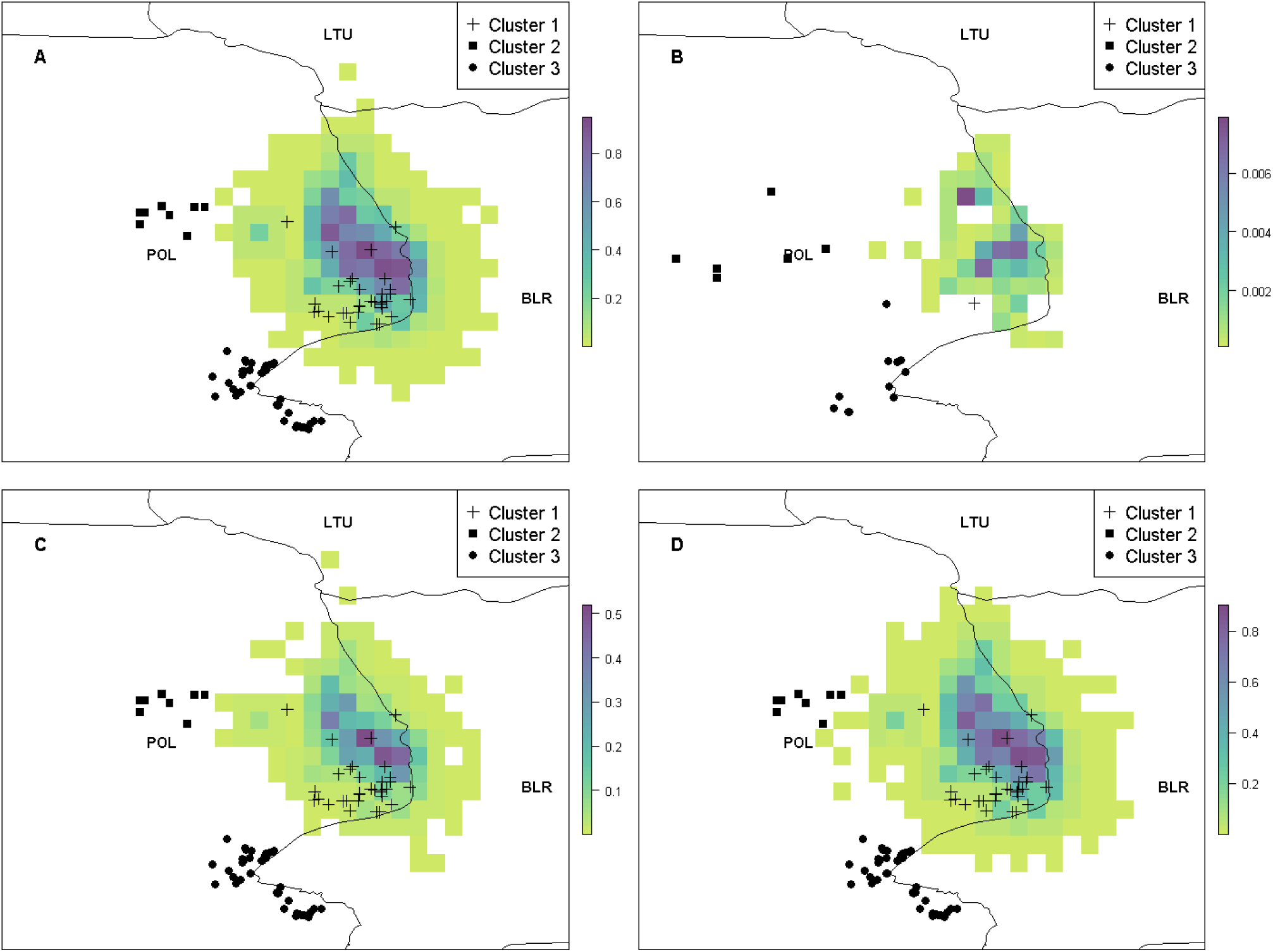
The probability of at least one new infection of ASF in boar and pigs in 2016 (A and B). In C and D, the probability of at least one infection of ASF in boar is split between transmission due to infected live boar and by contact with infected boar carcasses, respectively. In A, C and D boar cases in 2016 are plotted, while in B pig cases in 2016 are plotted. Each of the cases in 2016 are plotted with different shapes to represent which cluster the cases belong to according to the cluster analysis. Countries are represented by their ISO3 code.

We performed a cluster analysis of the 2016 cases of ASF, which indicated that there are 3 distinct clusters of cases that year (see Appendix B for methods and further results of the cluster analysis). This in itself suggests that the cases in the different clusters are unlikely to be caused by wild boar dispersion from the 2015 Poland cases as such spread would be expected to be more spatially homogenous. We indicate in Figure 4 which cluster all of the cases belong to. These are cluster 1 close to the cases in 2015 and then two separate clusters to the south (cluster 2) and to the west (cluster 3). The model indicates that it is highly unlikely (probability < 1e^-4^) that the cases of boar and pigs in the southerly or westerly clusters were due to wild boar movement from the 2015 Poland cases alone.

To investigate further, we consider when the reported cases of boar and pigs in 2016 occurred throughout the year (Figure 5). Pig cases only occurred in four months of the year whereas there were boar cases in nearly all 12 months. Figure 5 indicates that in both the southerly and the westerly clusters, the pig cases occurred first. For example, in the southerly cluster, most of the boar cases occurred in the 4^th^ quarter of the year, whilst all of the pig cases occurred in August and September, the 3^rd^ quarter. A similar story emerges in the westerly cluster. Our predictions indicate that these pig cases would not have been caused by wild boar movement as the probability of infection in pigs from boar movement is very low.

**Figure 5.**
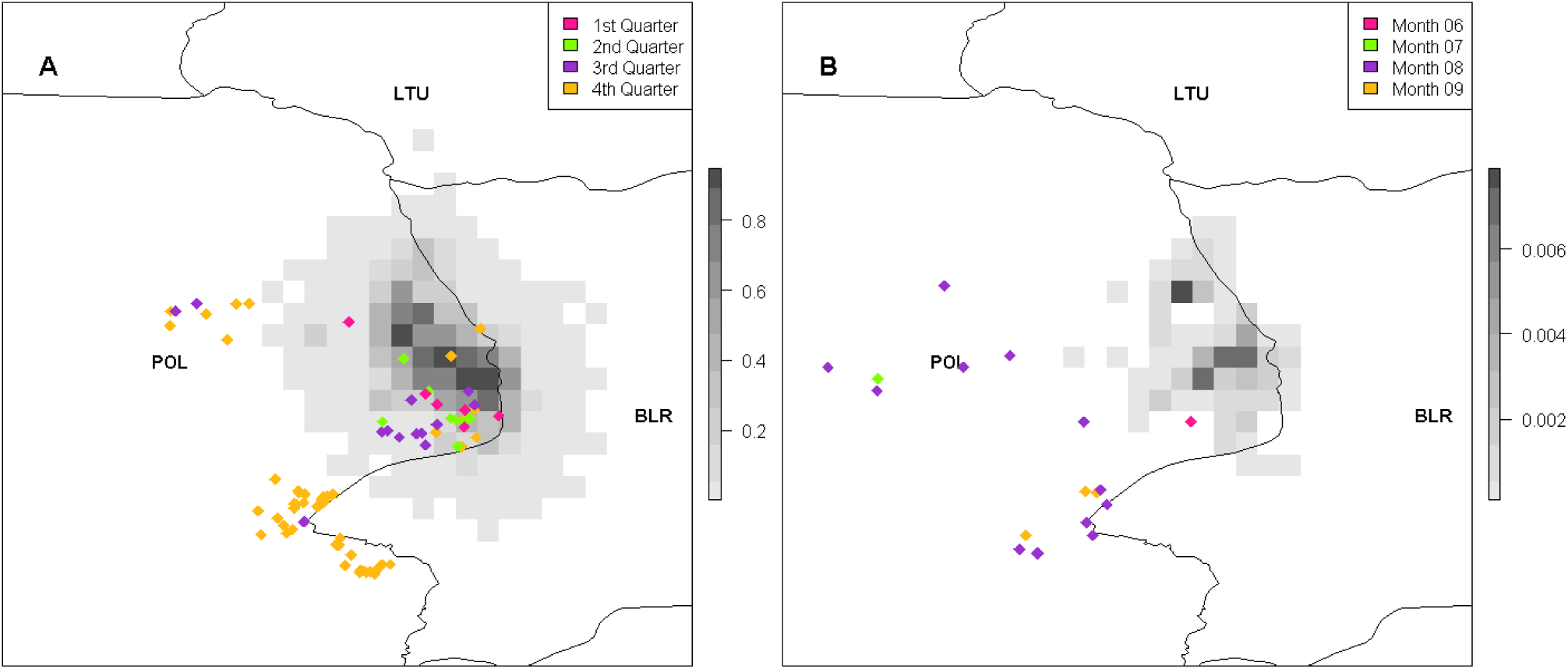
The probability of at least one infection of ASF in boar (A) and pigs (B) in 2016 is plotted in grey-scale while the reported cases of ASF in 2016 are plotted in colour depending on the time of year the case was reported. In A, boar cases are plotted based on the quarter of the year they occur: 1^st^ – Jan, Feb, Mar; 2^nd^ – Apr, May, June; 3^rd^ – July, Aug, Sept; 4^th^ – Oct, Nov, Dec. In B, pig cases are plotted according to the month in 2016 they occur. Countries are represented by their ISO3 code.

### Scenario Analyses

We compare the 10 control scenarios against the baseline scenario to assess their effectiveness using the severity and spread measures outlined above. We compute these metrics separately for boar (Figure 6). Full results of each of the scenarios are provided in Appendix C, for both boar and pigs.

**Figure 6.**
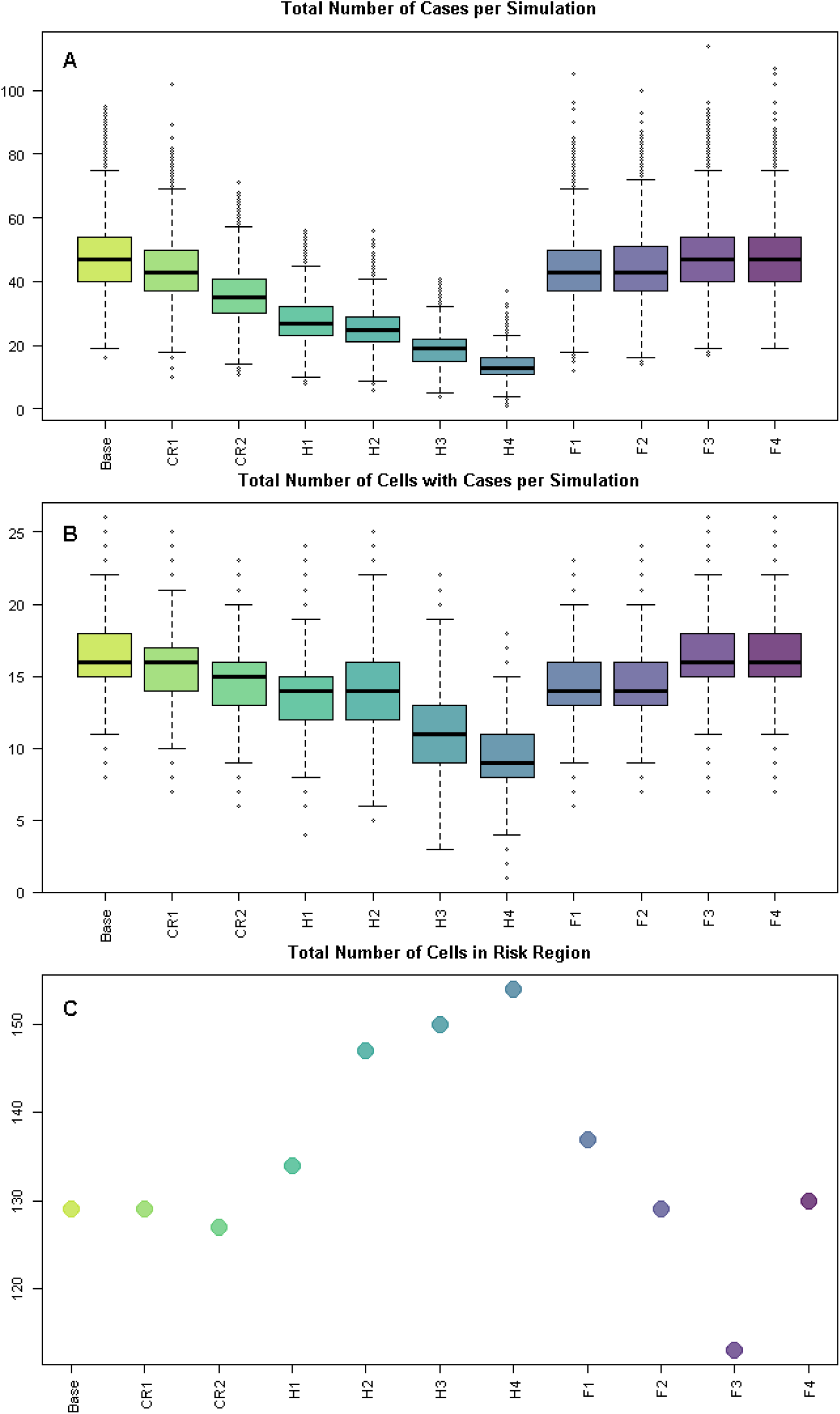
The baseline and scenario results are summarised using three metrics – the total number of new boar cases for each simulation; the total number of 100km^2^ cells which have at least one new boar case occurring for each simulation; and the total number of cells with at least one new case occurring over all simulations (i.e. the extent of the risk region). “Base” in the plot refers to the baseline 2015 results while each scenario is represented by a reference name as outlined in the methods section, where “CR” refers to the carcass removal scenarios, “H” to the hunting scenarios and “F” to the fencing scenarios.

With regard to reducing the total number of boar cases (Figure 6A), the hunting strategies appear to be the most successful, with H4, in which the boar population is reduced to ¼ of their original population size (in cells with cases in 2014) but 50% of boar undergo long-range dispersal, having the lowest number of cases per simulation. This strategy reduces the median number of cases per simulation by 72% from the baseline with H3 second at 60%. Perhaps surprisingly, H4 is a more successful strategy in reducing the total number of boar cases than H2 in which boar are also reduced to ¼ of their original population size but only 25% of boar undergo long-range dispersal. The reason for this is that there are more boar dispersing and therefore there is a greater likelihood that the boar will move outside of the original non-zero prevalence region. Normally, the area which had cases previously is the area where most cases will occur, either due to non-dispersing boar or boar dispersing to these cells as they are close by. Therefore, more boar dispersing leads to more boar leaving these cells. But there are many cells which the boar could move to, and hence each cell outside the non-zero prevalence area only receives a small number of boar and therefore the probability of a case occurring is low.

Similarly the number of cells which have a boar case per simulation is lowest for H4 followed by H3, with reduction by 44% and 31% respectively in the median number of cells infected per simulation. However, this does not mean that only 16 cells could be infected in total (the median for the baseline), but in each simulation on average 16 cells have infection occur within them. As can be seen from Figure 6C, the total number of cells that that have a non-zero probability of infection is highest for H4, with over 150 cells in the region at risk (an increase of 16% from the baseline). All of the hunting strategies lead to a bigger region of potential risk than the baseline scenario. This is due to the increase in the dispersal rate which leads to boar entering more cells further away from the original non-zero prevalence area, potentially causing infection in those cells with a low probability.

In comparison, the fencing strategy F3, in which a 95% successful fence of 20km width is built, is better for reducing the spread as less than 120 cells are in the risk region. This is a reduction of 14% from the baseline scenario. In the baseline scenario nearly all of the cases occur within 20km of the 2014 cases, thus the addition of a 20km fence does not reduce the total number of cases but it does reduce the possibility of infrequent cases occurring further away, provided the fence is effective. When the fence is not effective at 20km, scenario F4, the results are indistinguishable from the baseline scenario. On the other hand, a fence of width 10km is too narrow as it does not cover all the cells in which we have predicted non-zero prevalence based on 2014 cases, as can be seen in Appendix C. Thus, there are potentially infected boar outside the fence before the start of the simulation. In scenario F1, in which the fence is 95% effective, the dispersing infected boar outside of the fence are forced away from the fence, thereby spreading the disease further and resulting in a 6% increase in the region at risk. However, there is a reduction in total number of cases under this scenario, for a similar reason as above for hunting – the large number of cells to disperse to means fewer infected boar are entering each cell outside the fenced area and therefore there is less chance of a case occurring. Carcass removal (CR1, CR2) does not change the spread of the disease significantly as boar are still able to move the same amount, but it does reduce the chances of those dead boar infecting others and hence the total number of cases is reduced from the baseline, by 25% for CR2.

## Discussion

We have investigated the role of natural wild boar movement in the transmission of ASF within Poland during 2014-2016. Our model suggests that the role of wild boar movement may be limited to only very local spread. Even when cases were within a relatively short distance from the cases the previous year, such as the southerly and westerly clusters of cases in 2016, in none of our 10,000 simulations did we predict these cases due to wild boar movement. This is despite the fact that those clusters were 25 or 28km away from the 2015 cases, respectively, and boar in the model have the potential to move 50km. In comparison the cases from 2015 were much closer to the cases in 2014, with the maximum distance that a case in 2015 was from the nearest 2014 case of 9.163km. Similarly, the maximum distance that a case in cluster 1 in 2016 was from the nearest 2015 case is 16.889km. Overall, this indicates that on average, the spread of ASF has a moving boundary due to wild boar movement of less than 25km in a year. This is in line with statistical analysis of the cases in Europe, with estimates of spread at a rate of 2-5km/month (Chenais et al. 2019) and 1.5km/month (Podgórski and Śmietanka 2018). Our estimates, which are for spread by wild boar movement only, match the lower bounds of these statistical analyses. Overall, our findings regarding boar movement and ASF spread are in line with previous work, suggesting that wild boar movements have only a small role to play in ASF transmission (Podgórski and Śmietanka 2018) although this can depend on geographical location (Iglesias et al. 2018), and the fact that transmission by wild boar is driven principally by boar carcasses (Figure 3, Lange et al. (2018)).

The model suggests the westerly and southerly clusters of 2016 cases were not caused by wild boar movement from the 2015 Poland cases, and we corroborated this with the fact that cases in pig farms occurred first. Of course, it could be that cases were only found first in pigs and there was underlying infection in the boar population. However, as Poland was already aware of ASF in the country and were testing all boar carcasses found/hunted, it is unlikely that spread from the 2015 cases to these new clusters would have gone completely unnoticed, or that cases in the region between the clusters were not found, either before or after the pig cases. Therefore, we believe our model results reflect reality, and the conclusion has also been reached by the Polish authorities that the westerly cluster was caused first by human-mediated transmission to pigs and subsequent spillover to wild boar (pers. comm. General Veterinary Inspectorate) and supported by genetic tracking of ASF virus (Mazur-Panasiuk and Woźniakowski 2019). The main hypothesis for the original incursion in December 2013 in Poland is incursion from Belarus (Pejsak et al. 2014). The cases in 2016 in the southerly cluster are also very close to the Belarus border.

We have investigated the potential role that three control strategies, of varying effectiveness, could have on the transmission of the disease. Carcass removal overall had a positive effect on control of the disease, but the effect was minor. Hunting had the greatest potential to reduce the total number of cases but could also lead to the greatest region at risk. In contrast, fencing was successful or not depending on the width and the effectiveness of the fence. Fences which were only 50% effective did not reduce the severity or spread of the disease. Fences which had a width of 10km did not cover the total infected area and thus actually promoted spread since infected boar could only move further away from the fence if dispersing. Thus, only the fencing strategy of 20km and 95% effectiveness succeeded in reducing spread of the disease. However, it did not reduce the total number of cases as most cases occur within this 20km area. Choosing between these strategies, the risk manager has to weigh up whether it is more important to reduce the region at risk of the disease or reduce the total number of boar that could become infected from the original cases. Given the potential for spread to go unnoticed in an area for some time, it would seem that reducing the region at risk would be deemed most important. Lange and co-authors (Lange 2015, EFSA et al. 2017, Thulke and Lange 2017) have also considered the potential effects of control strategies separately, namely mass depopulation, a feeding ban and targeted hunting. These found that mass depopulation (provided it included prompt removal of carcasses) was the most successful strategy but the levels required was not feasible under conventional wildlife management practices and, furthermore, that measures were more effective if implemented preventively rather than in an area with infected cases. However, as already stated, many of the recent countries with new incursions have implemented a combined strategy of 3 control strategies (fencing, hunting and carcass removal) in different zones. We did not combine intervention strategies, as our focus was on the movement model rather than the control strategies, however, Lange et al. (2018) considered various strategies combining zoning, hunting, carcass removal and fencing. Similar to our control strategy of fences with 10km width, they found that fencing may be ineffective if based on locations of reported cases and not on locations of actual infected boar, but in general they found fencing to be a less successful strategy and instead focussed on combining intensive hunting with carcass removal. However, as also noted by Lange et al. (2018), further studies on boar ecology are required to understand how combined interventions may affect boar movement and group interactions, and therefore combining interventions within models is subject to high uncertainty.

Our model relies on data on many different aspects of the ASF situation, such as the reporting of cases, movement ecology, matrilineal group and territorial dynamics, transmission of disease and contact with carcasses. Added to this is the fact that these are all applied to a wild rather than domestic species, and thus undoubtedly there is less availability of or reliability in the data. For example, for many of our parameters, these are determined by studies on pigs as there are few experiments determining transmission and survival rates for wild boar. As the model is stochastic, many of our parameters incorporate natural variation which reduces our uncertainty in our ability to cover likely values of each parameter, even if there could still be uncertainty in the shape of the distribution. Most of the distributions for parameters are pert or uniform to incorporate our lack of detailed knowledge. Due to the high computational requirements needed to perform the movement section of the model stochastically, this part of the model is deterministic (i.e. there are no distributions regarding the number of steps a boar will take or the percentage of boar undergoing dispersal). While it is likely that different boar could have a wide range of dispersal distances, we assume that infected boar would not be able to move very far and thus the lack of variation in this parameter should not be so important. Similarly, there is a great deal of uncertainty over the proportion of boar carcasses that are found, and hence the under-reporting of boar cases. We did some visual inspections of scenarios involving parameters that were most uncertain to determine the effect variations in these parameters might have. While increasing the under-reporting factor or increasing the dispersal distance did increase the number of cases or the size of the risk region, as expected, they do not change our overall results, such as our conclusion that the southerly and westerly clusters in 2016 were not caused by wild boar movement from the 2015 Poland cases.

One aspect of the model which requires careful consideration is the role of secondary infections. Our model is only predicting the initial infections in new cells based on the estimated prevalence from reported cases. The model does not incorporate any other boar or pigs that could become infected due to the initial predicted boar cases. Therefore, we underestimate the total number of infected boar in an area. Furthermore, if all boar do not disperse around the same time each year, then it could be possible for an infected boar to disperse to a new cell, infect another boar in that cell, the new infected boar undergo its dispersal event after this (but still within the same year), and lead to infection in another boar or pig. This effectively stretches the size of the region at risk for one year and thus potentially indicates a faster wave-front of the disease spread. This issue is especially problematic under the hunting strategy as hunting disturbance could cause dispersal events to happen more frequently based upon when hunting occurs rather than the dispersal season. However, given the low probabilities involved in transmission, dispersal and movement distance, the increase to the risk region should be minimal as most cells outside of 20km already have very low probability of infection (< 2.5%, Figure *3*). One way to investigate the potential for increased spread by multiple dispersal events more rigorously would be to run our model on a shorter time scale than 1 year. Our model works for whatever time scale is most relevant, provided parameter rates are changed accordingly. For example, to run the model monthly, data on the percentage of boar dispersing each month would need to be supplied. This could include different timings of dispersal for young male boar dispersal compared to adult male and female dispersal. However it was not possible to do here due to the lack of data on when throughout the year dispersal events occur.

In all of the results, the model predicts the greatest risk in those cells which previously had cases (even in scenarios with high boar dispersal rates). This outcome is mostly occurring due to short distance movement as this always remains within the cell of origin. In reality, boar home-range could spread across multiple cells as boar are not restricted by our cell boundaries. Further, in the model boar may choose to disperse to cells quite close by and therefore could move into cells that already had cases. Perhaps, in reality, boar would be more directed in their movement away from their original location (Morelle et al. 2015). Another potential explanation is that the model underestimates the effect of the disease within the boar population and specifically on the rates of boar dispersal. It may be that the dispersal rate increased due to presence of the disease or disturbance from human intervention, whether carcass removal or other control strategies. Lastly, it could be that there are boar becoming infected but are recovering from the disease, and are therefore not reported as cases since cases are predominantly found via boar carcasses rather than active testing of the boar population. However, this seems unlikely given the current knowledge of the highly virulent nature of the disease, even when boar and pigs are infected with low doses (Pietschmann et al. 2015).

While we have investigated the role that wild boar movement has in local transmission of ASF, other transmission pathways can also be important, such as movement of pigs from farm to farm, contaminated cars, shoes or clothing, or transport/removal of boar carcasses if the necessary biosecurity is not followed. We have not assessed the risk of ASF in new areas in Poland due to these transmission pathways as the focus was on understanding the role of wild boar movement. However, this model framework for transmission of ASF by wild boar movement can now be combined with other pathways in order to compute the risk of infection of ASF on a broader scale through many routes. The fine spatial scale of the model favours results that can be used to target surveillance activities by highlighting potential areas of high risk of infection that have yet to report cases. For example, as seen in Figure 2 and Figure 4, there was a high risk of cases in wild boar in Belarus on the border with Poland during both 2015 and 2016 and hence the model could have been used to indicate locations best to test for ASF in Belarus. Although the model was shown using a case study for Poland in 2014-2016, it is applicable across the whole of Europe, provided data on boar abundance and habitat suitability is available. Furthermore, the model can be run prospectively, using up-to-date data on locations of reported cases. Combining the ability to run the model on real-time data and for the whole of Europe at a fine spatial scale, the model has the potential to aid in ASF hotspot identification, and consequently in developing risk-based surveillance plans, a necessity if ASF will be successfully controlled across Europe and worldwide.

## Acknowledgements

This project is part of COMPARE (Collaborative management platform for detection and analysis of (re-) emerging and foodborne outbreaks in Europe) which received funding from the European Union’s Horizon 2020 research and innovation programme under grant agreement No 643476. T. Podgórski was financially supported by the National Science Centre, Poland (grant number 2014/15/B/NZ9/01933).

