## Supplementary Information for "Predicting spread and effective control measures for African swine fever– should we blame the boars?"

#### Appendix A: Implementation of the Scenario Analysis

We perform different scenario analyses to address the potential control strategies that could have been implemented in Poland to assess their effectiveness when wild boar alone is responsible for transmission. We assume that the control strategy is implemented in 2014 and compute a new risk for 2015 including the potentially positive and negative effects of the control strategy. We then compare our new results against the baseline results for 2015.

The scenario analyses we perform are:

A. Increase in carcass removal:
   This affects the probability of a carcass being found and removed; in the baseline model 1 in 4 carcasses are removed
   - CR1. 1 in 3 carcasses are found and removed
   - CR2. 1 in 2 carcasses are found and removed

B. Hunting:
   This affects both the number of boar in the area and the probability that boar will move due to disturbance
   - H1. Reduction of boar population to half original size and 25% of remaining boars disperse
   - H2. Reduction of boar population to quarter of original size and 25% of remaining boars disperse
   - H3. Reduction of boar population to half original size and 50% of remaining boars disperse
   - H4. Reduction of boar population to quarter of original size and 50% of remaining boars disperse

C. Fencing:
   The implementation of fencing around the current cases is determined by a buffer width and the permeability of the fence in successfully stopping boar moving outside the buffer
   - F1. Fence of 10km buffer and 95% successful
   - F2. Fence of 10km buffer and 50% successful
   - F3. Fence of 20km buffer and 95% successful
   - F4. Fence of 20km buffer and 50% successful

The implementation of the carcass removal strategy is performed simply by increasing the parameter "probability that a carcass will be removed" from ¼ to 1/3 or ½. Similarly, adding the control strategy of hunting is relatively straightforward. It involves changing the parameter "proportion of boar performing long range movement" and reducing the susceptible population, $S(c)$, by the relevant amount in each cell where cases were reported. We do not adjust the population size in all cells which we estimate have non-zero prevalence (as we have used a smoothing method to estimate prevalence), only those cells which had reported cases.

The implementation of fencing scenarios is a little more complicated. For both fence widths (10km and 20km), we draw a buffer of the relevant size around the reported cases in 2014 (Figure 1). As can be seen from Figure 1, the 20km fence does not go through any cells with non-zero prevalence, because our smoothing method only extends to direct neighbours and our cells are 10x10km. Therefore, for the 20km case, we set the suitability score $h(c)$ for all cells $c$ which contain the fence based upon the success of the fence – if the fence is 95% successful, we reduce the suitability score in all cells with a fence to 5% of their original score. Thus, the revised suitability score $h^*(c)$ is given by

$$h^*(c) = h(c) * p_{Fence},$$

where $c$ represents all cells the fence goes through, $h(c)$ is the original suitability score of cell $c$ and $p_{Fence}$ is the probability that a boar will get through the fence. We then calculate the movement model as before, with the assumption that any boar which enter a cell with the fence in it have effectively crossed the fence.

For the fence of 10km wide, we need to be more careful, as there are non-zero prevalence cells which have the fence going through them. For the cells that are completely inside the fence, we use the same method as above, adjusting the suitability score in the cells with fence running through them. However, for the cells which have the fence going through them, we first need to determine if the infected boar in that cell are inside or outside the fenced zone at the start of the simulation. To do this, we split the cell into two parts; inside and outside the fence. The number of boar in each part is assumed to be proportional to the area of the cell in each part. For those boar determined to be inside the fence, we adjust the suitability score of the neighbouring cells outside the fence using the success score of the fence, as before. We also set the number of boar in the fence cells that these boar can have contact with to the size of the boar population inside the fence, as we assume they will not be able to contact the boar in the same cell which are outside the fence. Hence a boar in that cell can only interact with the boar population inside the fence, unless it successfully moves to a cell outside the fence buffer (which now has a lower probability due to the reduction in the suitability score). Lastly, for the boar outside the fence but in a cell with a fence, we reverse the above calculation. We set the boar population of these fenced cells to be the proportion outside the fence. We then reduce the suitability of the cells which are neighbouring cells *inside* the fence. Therefore, if a boar successfully moves into those cells, we assume that the boar has effectively moved into the fenced area.

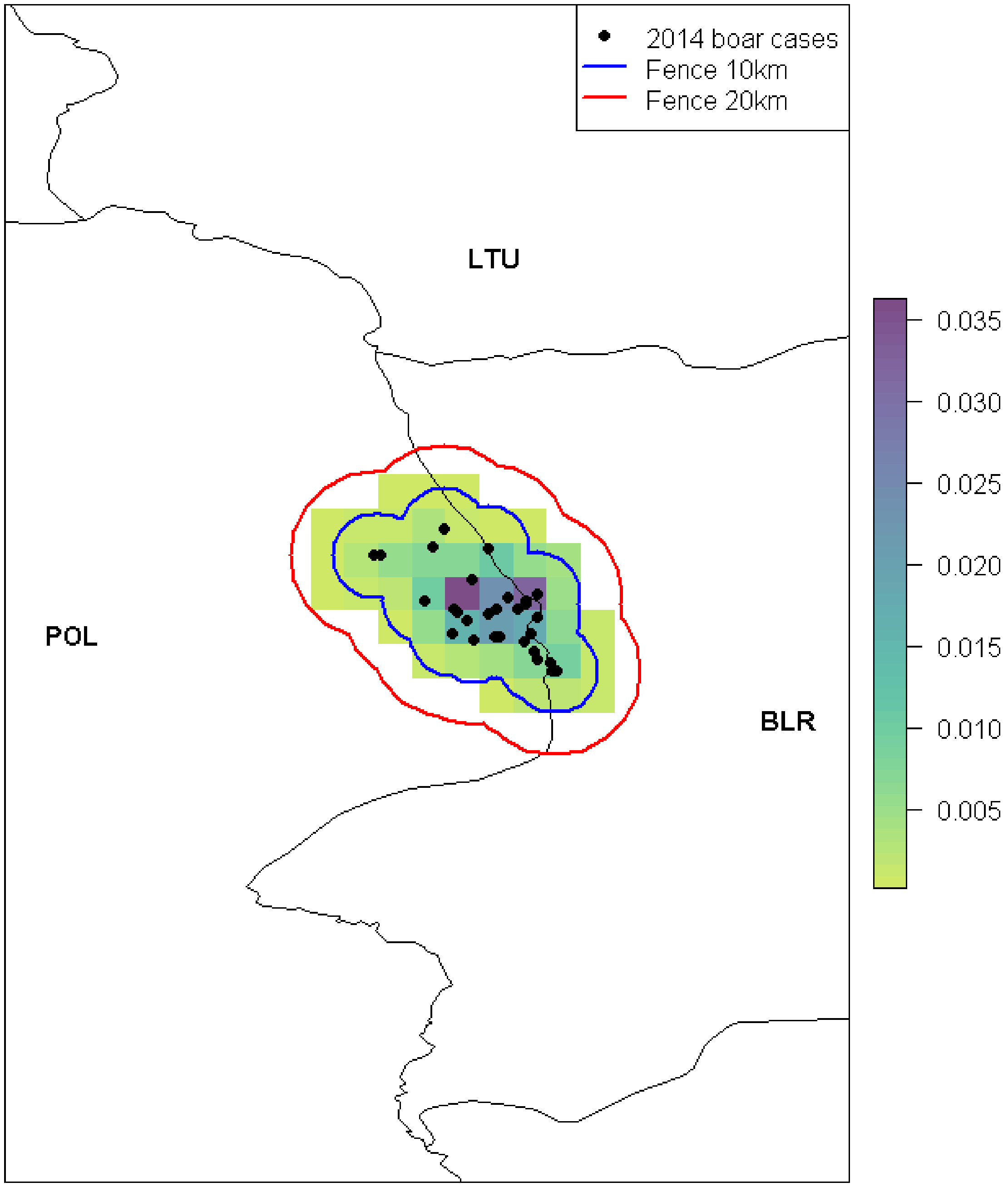


Figure 1  The prevalence of ASF in 2014, estimated using the reported cases in 2014, is plotted at a 100km$^2$ cell level. A buffer of 10km (blue) and 20km (red) is plotted around the 2014 cases indicating the fencing in the four fencing control scenarios. Black circles represent the reported cases of ASF in wild boar in 2014.

### Appendix B: Additional Baseline Results

**Probability of new infections in 2015 given 2014 reported cases**

A plot of the probability of cases in boar, depending on whether the transmission occurred via live boar or boar carcasses is provided in Figure 2. The highest probability in any cell that a case will occur via live boar transmission is 0.63 whereas it is 0.94 for transmission via boar carcasses. Furthermore, there is a wider region of risk via carcass transmission compared to live boar transmission.

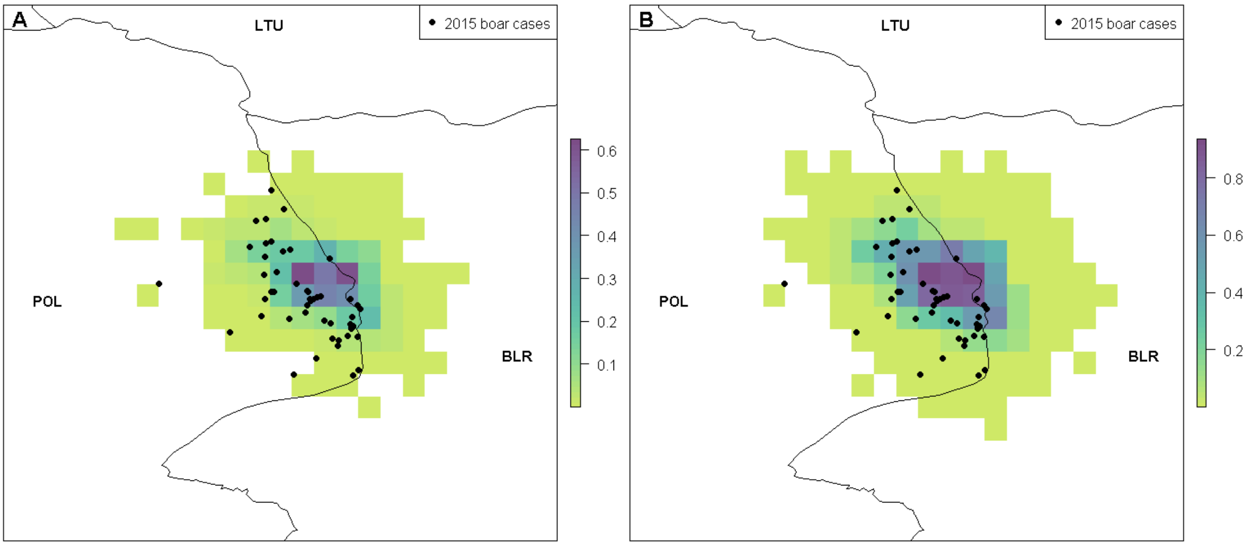


Figure 2 The probability of at least one infection in boar due to transmission via live boar contact (A) or contact with dead boar carcasses (B) over one year. Black circles represent reported cases of ASF in boar and pigs in 2015. Countries are indicated by their ISO3 code.

**Probability of new infections in 2016 given 2015 reported cases: Cluster Analysis**

We performed a kmeans cluster analysis on the 2016 reported cases using the function kmeans in R (R Core Team 2017), a common method of performing cluster analysis in machine learning. This function aims to split data into different clusters based on their similarity, in this case based on their geographical similarity. It assigns cases to distinct clusters in order to reduce within-cluster variance. To use this method the number of clusters you want the cases to be split into needs to be assigned in advance. We tested 1 to 5 clusters and, using the “elbow method” on the within sum of squares, we discover that the best number of clusters is 3 as this reduces the within-cluster sum of squares without overfitting. See Figure 3.

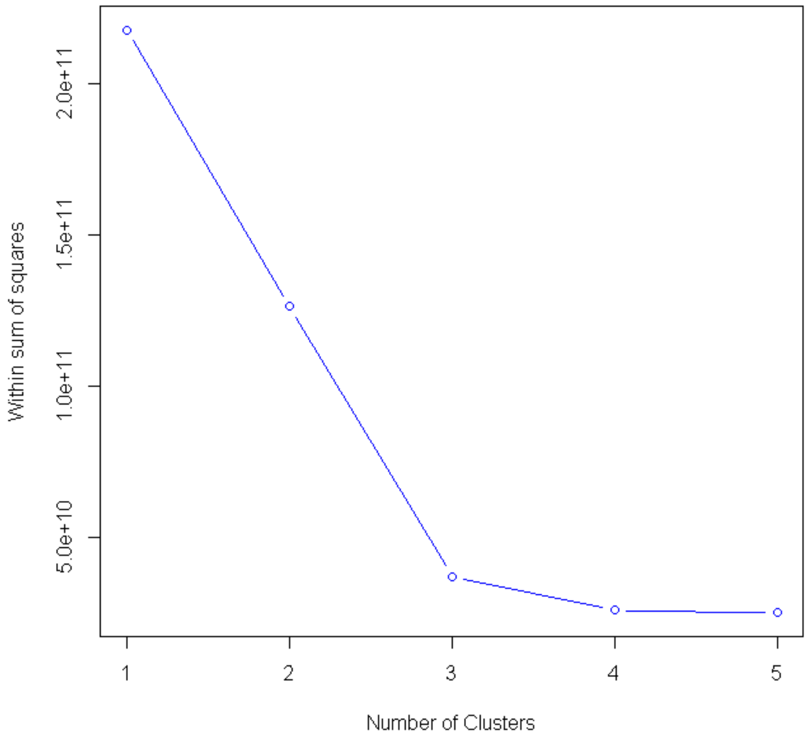


Figure 3 The within-cluster sum of squares is plotted against the number of clusters to determine which number of clusters is most optimal. The optimal value, by the "elbow method" is where the bend in the graph appears, in this case at 3.

**Probability of new infections in 2016 given 2015 reported cases: Quantiles**

We present 2.5%, 50% and 97.5% quantiles for the predicted number of new cases in wild boar in 2016 using the reported cases in 2015 to estimate prevalence (Figure 4). The number of new cases is split between whether the case occurs due to transmission via live boars or via boar carcasses.

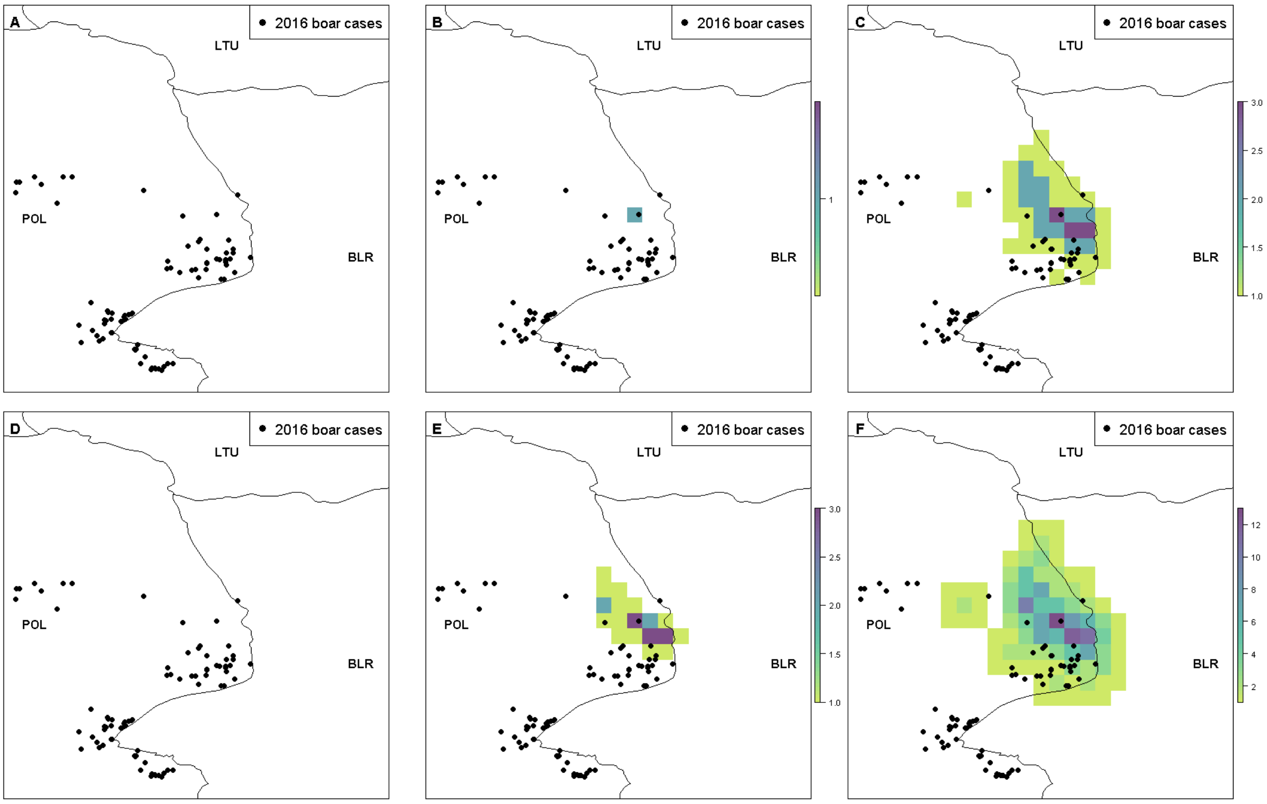


Figure 4 The 2.5%, 50% and 97.5% quantiles of the number of new infections in boar in 2016 due to transmission by live boar (A, B, C respectively) and by contact with boar carcasses (D, E, F respectively). Black circles indicate reported cases of boar in 2016. Countries are indicated by their ISO3 code.

### Appendix C: Full Results of Scenario Analysis

#### Comparison of Scenarios for Pig Cases

We include the comparison of the scenarios using pig cases in Figure 5.

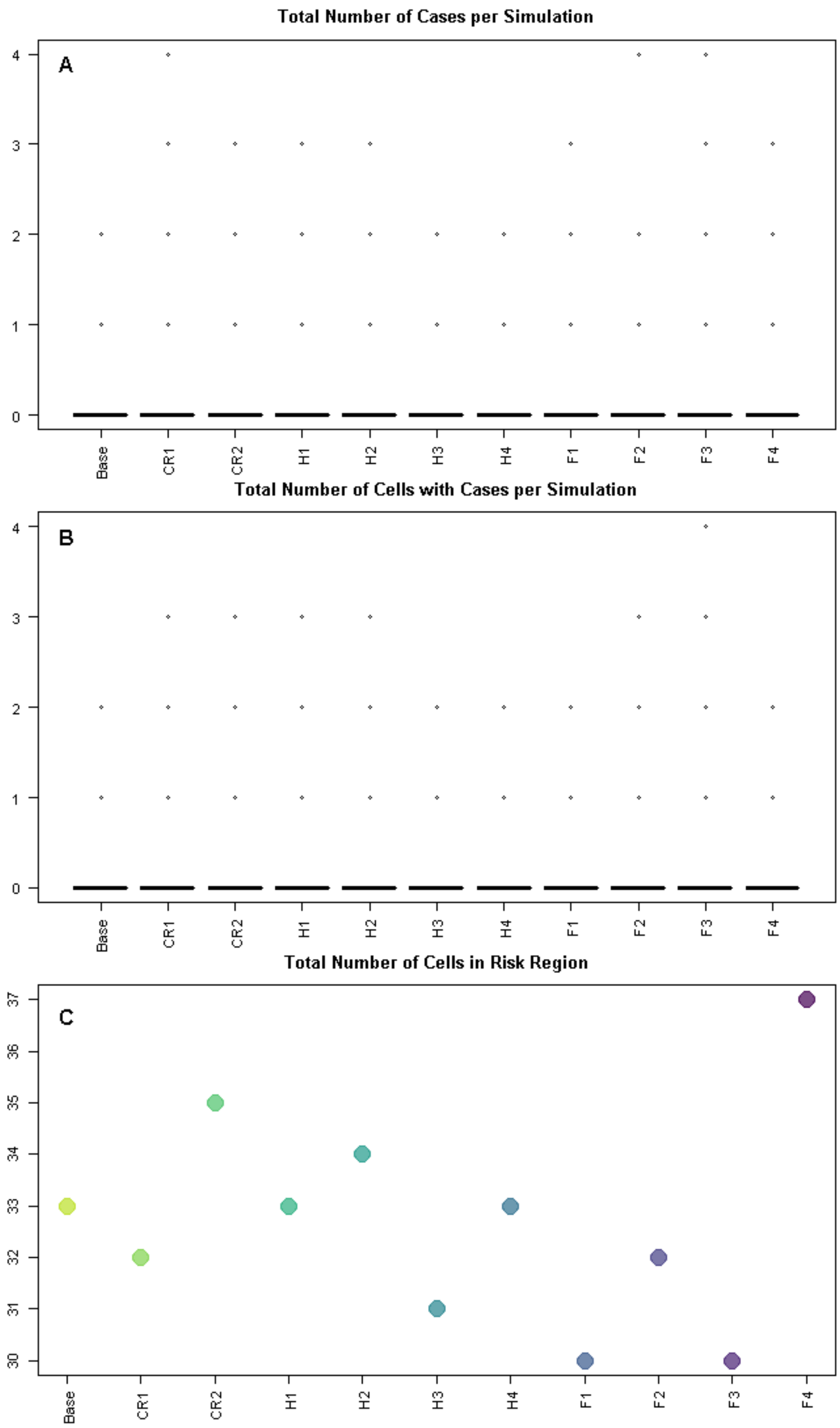


Figure 5 The baseline and scenario results are summarised using three metrics for pig cases – the total number of new pig cases for each simulation; the total number of 100km$^2$ cells which have at least one new pig case occurring for each simulation; and the total number of cells with at least one new case occurring over all simulations (i.e. the extent of the risk region). "Base" in the plot refers to the baseline 2014-2015 results while each scenario is represented by a reference name as outlined in the methods section, where

“CR” refers to the carcass removal scenarios, “H” for the hunting scenarios and “F” for the fencing scenarios.

As the probability of pig cases occurring is so low, the differences between the scenarios is not very apparent and is most likely driven by stochastic differences.

#### Scenario Results for Boar Infection

We show the spatial maps for the probability of at least one case in boar for each of the scenarios together (Figure 6). This indicates the increased size of the risk region for the hunting strategies H3 and H4 as well as the reduction in cases (since the maximum probability of cases reduces from 0.97 in the baseline to 0.68 for H4). Also, it illustrates the reduction in the risk region for the fencing strategies, especially F3.

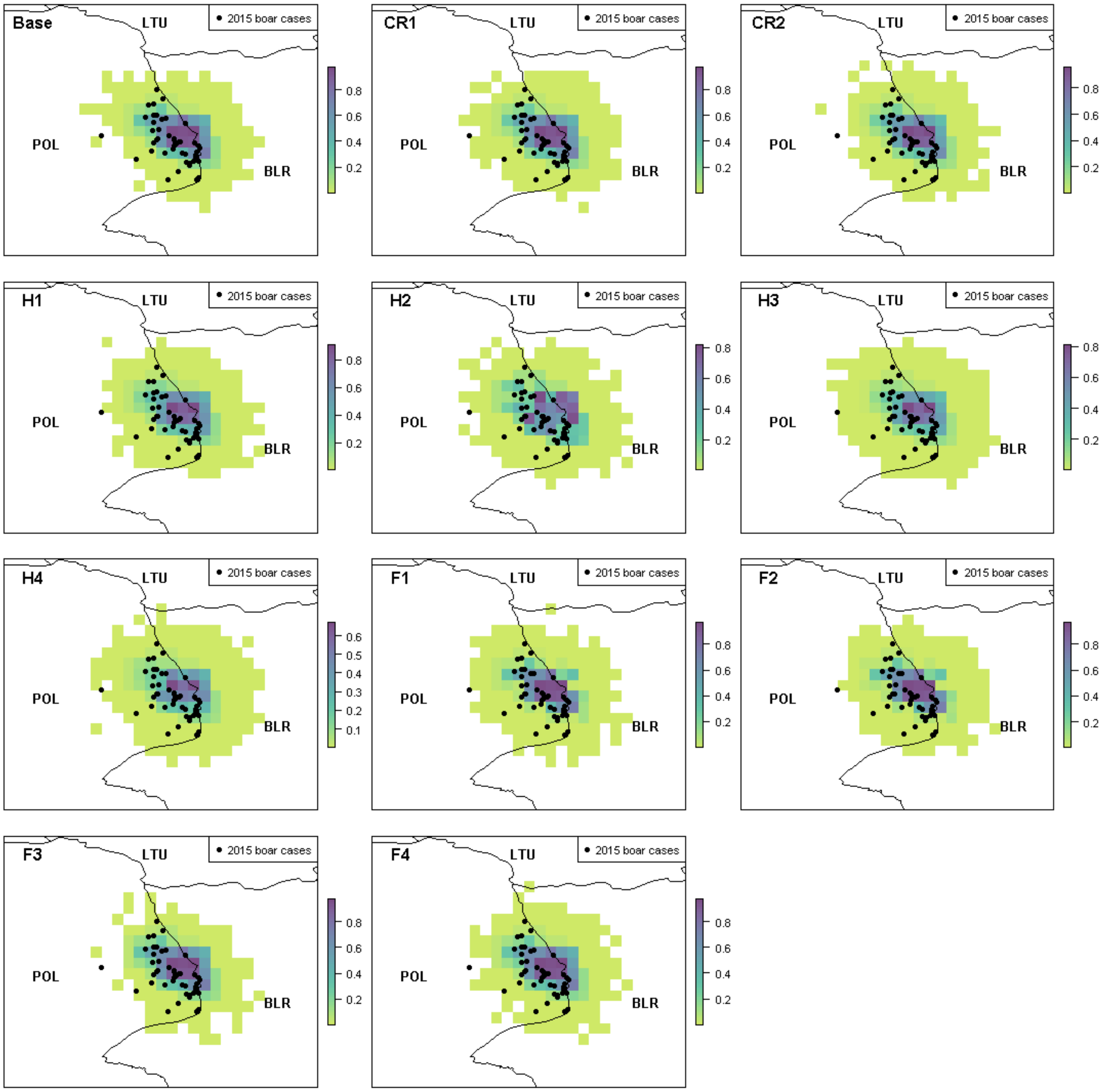


Figure 6 The probability of at least one infection in boar is plotted at a 100km$^2$ cell level in 2015 under the baseline scenario and the 10 control strategies. “Base” in the plot refers to the baseline 2014-2015 results while each scenario is represented by a reference name as outlined in the methods section, where “CR”

refers to the carcass removal scenarios, “H” for the hunting scenarios and “F” for the fencing scenarios. Black circles indicate reported cases of boar in 2015. Countries are indicated by their ISO3 code.

#### Scenario Results for Pig Infection

We show the spatial maps for the probability of at least one case in pigs for each of the scenarios together (Figure 7), indicating that the scenarios do not make much difference to the spatial probability of at least one case in pigs.

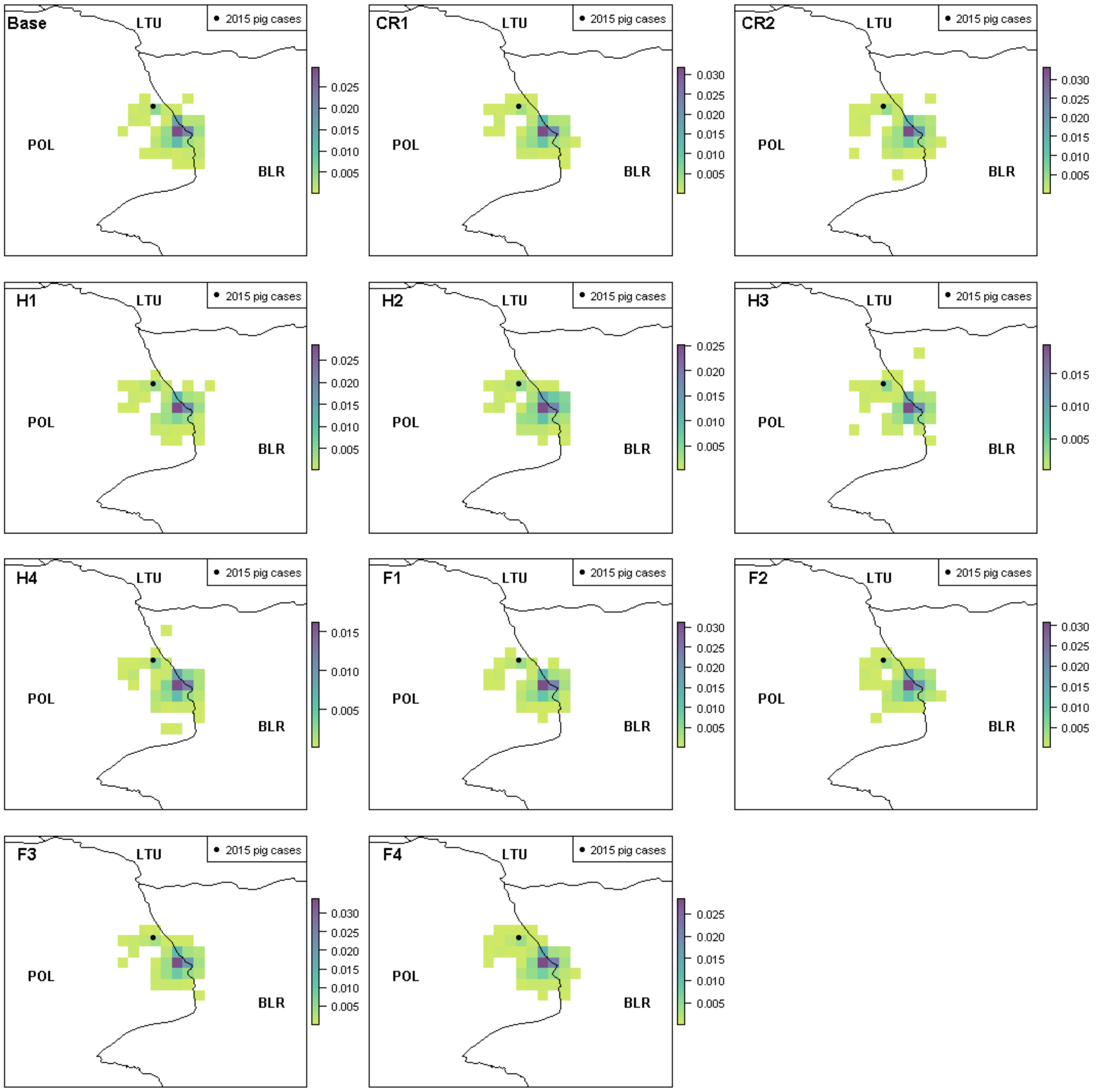


Figure 7 The probability of at least one infection in pigs is plotted at a 100km$^2$ cell level in 2015 under the baseline scenario and the 10 control strategies. “Base” in the plot refers to the baseline 2014-2015 results while each scenario is represented by a reference name as outlined in the methods section, where “CR” refers to the carcass removal scenarios, “H” for the hunting scenarios and “F” for the fencing scenarios. Black circles indicate reported cases of pigs in 2015. Countries are indicated by their ISO3 code.
